# Evolutionary remodeling of non-canonical ORF translation in mammals

**DOI:** 10.1101/2025.09.16.676541

**Authors:** Yue Chang, Tianyu Lei, Feng Zhou, Jiawen Jiang, Yu Huang, Ziyang Zhu, Hong Zhang

## Abstract

Non-canonical open reading frames (ncORFs) are pervasive within transcripts annotated as “non-coding” or “untranslated regions” of mRNAs, yet their landscape under normal physiological conditions remains to be fully resolved, particularly outside humans. Here we applied a stringent and standardized pipeline to hundreds of high-quality ribosome profiling libraries from normal mammalian tissues and cell types, identifying 11,623 human and 16,485 mouse ncORFs. Evolutionary analyses revealed that thousands of ncORFs are subject to coding constraint and exhibit lineage-specific conservation, underscoring their functional potential. Ancient ncORFs are preferentially highly translated, broadly expressed, and enriched for lineage-specific conservation. Co-expression patterns further indicate that many ncORFs, especially ancient ones, are cotranslated with canonical coding sequences, consistent with functions mediated through protein–protein interactions. Together, these findings establish a comprehensive atlas of mammalian ncORFs and provide fundamental insights into their evolutionary dynamics and functional integration within the proteome.

## Introduction

The systematic identification of functional genomic elements represents a foundational objective in the field of genomics. Among these elements, non-canonical open reading frames (ncORFs)—which are open reading frames (ORFs) located within long non-coding RNAs (lncRNAs) or untranslated regions (UTRs) of mRNAs—have attracted considerable attention in recent years^1–5^. While certain ncORF subtypes, such as upstream ORFs (uORFs) located in 5’ UTRs and downstream ORFs (dORFs) in 3’ UTRs, have been extensively characterized for their ability to regulate mRNA translation and stability in *cis*, a growing body of evidence demonstrates that ncORFs can also give rise to functional proteins that engage in diverse cellular pathways^1,2,6,7^. As functional ncORFs continue to be discovered across multiple eukaryotes^8–17^, systematic elucidation of their composition and function promises to uncover an additional layer of regulatory complexity and to refine our conception of the so-called “non-coding” genome.

Although ncORFs may play important biological roles, the number of translated ncORFs in the human genome remains uncertain. Detection by mass spectrometry (MS)-based proteomics is challenging, likely due to their short lengths and low abundances^18^. Ribosome profiling (Ribo-Seq) offers greater sensitivity for identifying translation events^18,19^, but downstream computational pipelines often produce highly divergent ncORF sets ^20^. Consequently, published estimates of ncORF abundance span several orders of magnitude—ranging from thousands to millions ^15–22^. To establish a standardized reference, the GENCODE consortium integrated data from seven distinct studies to produce a catalog of 7,264 human ncORFs (hereafter GENCODE ncORFs)^2^; however, the heterogeneity of sample sources and prediction algorithms likely introduced systematic bias. This catalog is also restricted to AUG-initiating ORFs, thereby overlooking those with near-cognate start codons. A later effort generated a high-resolution human translatome map and identified 7,767 ncORFs with a unified pipeline^22^. Nevertheless, this conservative annotation disproportionately under-represents coding sequence (CDS)-overlapping ncORFs. Indeed, later estimation suggests that between 10,500 to 22,500 ncORFs should exist in the human genome^18^. Furthermore, many ncORFs were reported from studies of cancer cells^1,13,23,24^, where extensive transcriptome reprogramming alters the translational landscape^25,26^. While paired healthy tissues were also analyzed in these studies, the landscape of ncORFs translated under normal physiological conditions is not yet fully resolved.

While a limited subset of ncORFs has been shown to encode signaling molecules, cofactors, membrane proteins, or subunits of large protein complexes^1,27,28^, functions of most ncORF-encoded proteins (ncEPs) are largely uncharacterized. Advances in high-throughput screen technologies have enabled the systematic interrogation of functional ncEPs^23,28–32^. Nevertheless, there is minimal concordance among hits from different studies^3^. Evolutionary analyses can offer insights into potential functionality of these translation products and mechanisms of origination^33–35^, yet how ncORF translation has been remodeled to integrate into proteome since their evolutionary origins is still poorly understood. Consequently, the extent to which translated ncORFs encode functional proteins and the evolutionary dynamics of their integration into the existing proteome are largely unresolved. Moreover, because most ncORF studies have focused on humans, it is unclear whether ncORFs in other species share similar sequence features and expression patterns.

To address these gaps, we generated a standardized annotation of translated ncORFs by uniformly reprocessing ~400 high-quality ribosome profiling libraries from normal mammalian tissues and cell types. This stringent approach yielded 11,623 human and 16,485 mouse ncORFs, consistently recovered across diverse samples and prediction strategies. Comparative and functional genomic analyses revealed that thousands of these ncORFs are under evolutionary constraint and exhibit distinct translation profiles. The resulting dataset provides a robust, comprehensive resource that will facilitate future investigations into the biological roles of ncORFs across cellular and organismal contexts.

## Results

### Annotation of ncORFs translated in normal mammalian cells

We developed a standardized pipeline to systematically annotate ncORFs translated in normal human and mouse cells (Fig. 1A). We first curated publicly available Ribo-Seq libraries, excluding those from pathological or genetically modified samples. Since both library size and the proportion of in-frame ribosome-protected fragments (RPFs) affect the sensitivity and accuracy of ncORF prediction^20^, we filtered out libraries containing fewer than 5 million RPFs or with in-frame rates below 60%. After this quality control step, we retained 174 human and 209 mouse high-quality Ribo-Seq libraries, spanning a range of tissues and cell types (Fig. S1A; Table S1). We predicted translated ORFs that begin with NUG (where N stands for any nucleotide) start codons and encode at least five amino acids using three tools—PRICE^36^, RiboCode^37^ and Ribo-TISH^38^—which we previously benchmarked for optimal performance^20^. After excluding annotated CDSs and their extension or truncation variants arising from alternative initiation sites, we obtained a preliminary list of ncORFs in each library.

**Figure 1.**
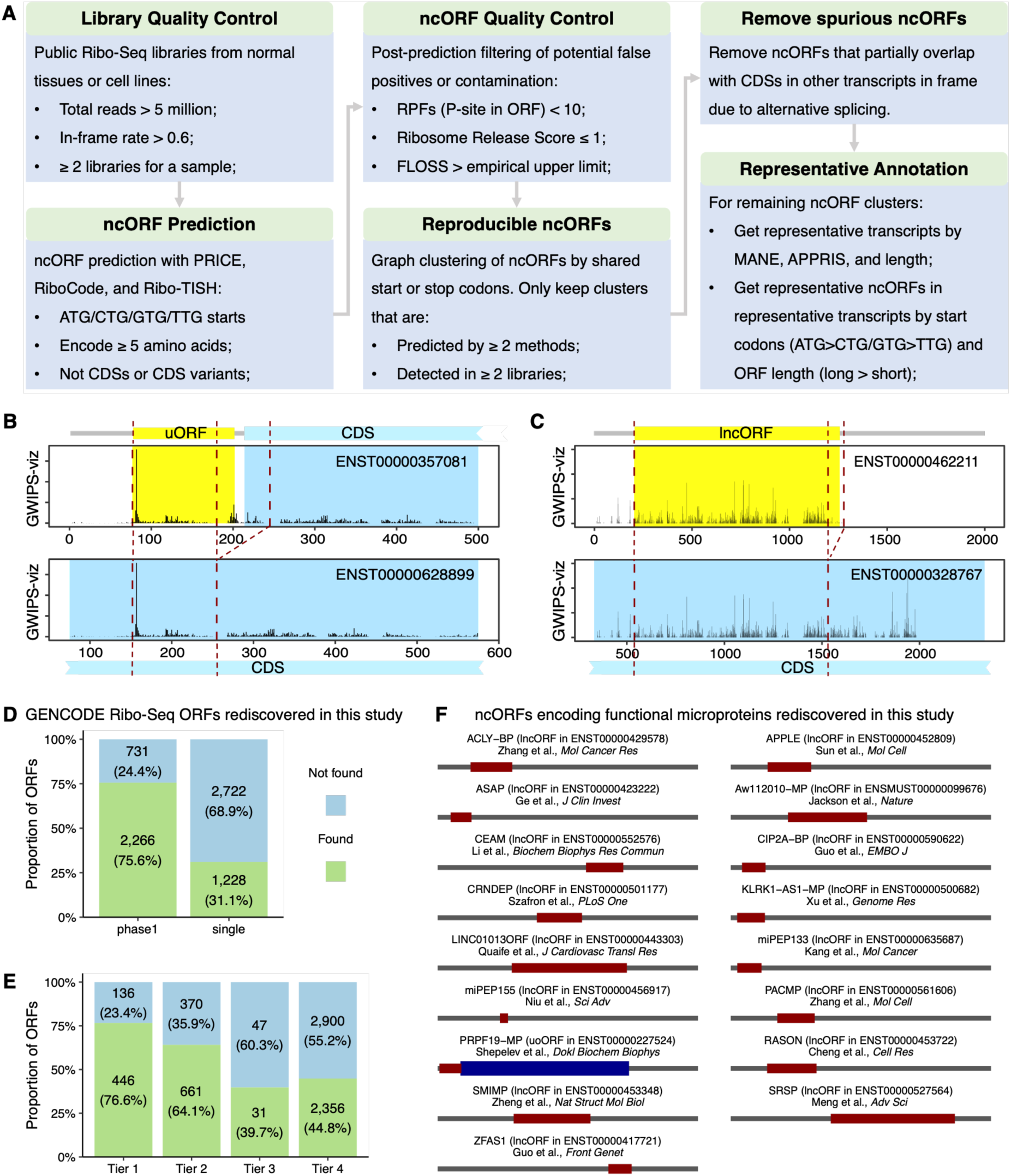
Annotation of ncORFs in humans and mice. (**A**) Overview of the integrative pipeline for ncORF annotation in human and mouse genomes. (**B-C**) RPF coverage plot for a representative uORF (B) and a lncORF (C), visualized using the Ribo-Seq signal track from GWIPS-viz^72^. ncORFs were shown in yellow and CDSs in cyan. Dashed lines indicate identical genomic regions shared across transcript isoforms. (**D-E**) Proportion of GENCODE ncORFs rediscovered in this study. GENCODE ncORFs were stratified either by reproducibility across independent studies (D) or by tiered translation evidence strength (E). (**F**) Known ncORFs that were experimentally characterized in earlier studies and independently rediscovered in this study from humans and mice. CDS, coding sequence; FLOSS, fragment length organization similarity score; MANE, matched annotation from NCBI and EBI; ncORF, non-canonical open reading frame; ORF, open reading frame; RPF, ribosome-protected fragment.

To generate reliable ncORF annotations, we applied stringent filtering and integration criteria. We removed potential false positives caused by non-ribosomal contamination or translational noise based on metrics including Fragment Length Organization Similarity Score (FLOSS)^39^, Ribosomal Release Score (RRS)^40,41^ and the number of RPFs within an ORF by P-site. We then aggregated the remaining ORFs to identify those consistently detected across multiple methods and libraries in human and mouse datasets, respectively (Fig. 1A). In many cases, two ORFs predicted from different libraries or methods may share the same start or stop codon due to the alternative translation initiation or splicing, resulting in groups of overlapping ncORFs. For example, 11.1% (805) of GENCODE ncORFs overlap with at least one other ORF, forming 329 overlapping groups (Table S2). This overlap complicates the identification of reproducible ORFs, as ORFs within a group may represent distinct proteoforms of a protein. To address this issue, we collapsed each group of ncORFs sharing the same start or stop codon into a single cluster, and retained only those clusters supported by at least two independent methods and observed in more than one library. For each retained cluster, we selected a representative ORF by prioritizing transcript annotation^42,43^, start codon type, and ORF length to facilitate downstream analyses (Fig. 1A).

This approach yielded 12,469 translated ncORFs in humans and 16,960 in mice. However, manual inspection revealed a subset of spurious ncORFs that are not truncation or extension variants of annotated CDSs, but instead partially overlap with annotated CDSs from alternative transcript isoforms in frame (Figs. 1B & C). For instance, in gene *AGTPBP1*, the CDS of transcript *ENST00000357081* begins downstream of *ENST00000628899* CDS due to the inclusion of an alternative exon. As a result, the 5’ UTR of *ENST00000357081* appears to contain a translated uORF (Fig. 1B), which overlaps in-frame with the CDS of *ENST00000628899* and may be mistakenly identified as a translated ncORF owing to active translation of the latter. A similar case is seen with the lncRNA ORF (lncORF) in *ENST00000462211*, which overlaps in-frame with the CDS of *ENST00000328767* from the *TMTC4* gene and may thus be erroneously identified as a translated ncORF (Fig. 1C). To avoid such artifacts, we implemented an additional filtering step to exclude these overlapping cases and left with 11,623 and 16,485 high-confidence ncORFs in humans and mice, respectively (Table S3). Notably, while more than half ncORFs were detected in two to five libraries, 22.3–26.3% were identified in more than 10 libraries, indicating a substantial subset with robust and recurrent translation signals suitable for future experimental validation (Fig. S1B).

### Annotated ncORFs are supported by independent evidence

To assess whether the annotated ncORFs are supported by independent evidence, we compared them with a previously compiled catalog of ncORFs identified using mass spectrometry (MS)-based proteomic data^20^. For the 3,494 MS-supported ncORFs in humans and 873 in mice, 44.5% and 67.6%, respectively, were also predicted as translated ncORFs in this study (Fig. S2). We further evaluated our annotation against the GENCODE ncORFs^2^, a widely used community resource. Of the 7,264 GENCODE ncORFs, 6,974 are located in fully annotated transcripts that were included in our analysis. Among them, 2,997 were reported by at least two independent studies (hereafter “phase 1”), while the remaining were only reported in a single study (“single”)^2^. We found that 75.6% of phase 1 ncORFs were independently recovered in our annotation, significantly more than the single-study ncORFs (Fig. 1D; *P* = 6.4×10^−307^, Fisher’s exact test).

A recent study conducted a comprehensive pan-proteome analysis to provide high-confidence peptide evidence for GENCODE ncORFs^44^. In this study, ncORFs were categorized into four tiers: tier 1 and 2 ncORFs had support from both MS and Ribo-Seq data; tier 3 had MS evidence only; and tier 4 had Ribo-Seq evidence only^45^. Notably, our annotation was significantly enriched for tier 1 or 2 ncORFs (*P* = 2.0×10^−64^, Fisher’s exact test), with 446 and 661 of them detected, respectively (Fig. 1E). Combining MS-supported ncORFs from our previous compilation and those from GENCODE ncORFs, 2,088 (18.0%) human ncORFs identified in this study were supported by MS evidence. Despite this overlap, the majority (65.2%) of human ncORFs in our annotation are not included in either of the two catalogs. This highlights both the complexity of the non-canonical translatome and the critical role of tissue diversity in comprehensive ncORF annotation.

To further validate our findings, we manually curated ncORFs with known functions from the literature. We identified 17 ncORFs in our annotation that have been experimentally confirmed to be translated—either through reporter assays or antibody detection—and shown to produce functional proteins (Fig. 1F; Table S4). Notably, 16 of these validated ncORFs are from humans, and only 12 are included in the GENCODE catalog. This analysis further underscores the accuracy and functional relevance of our annotations. Together, these results indicate that our annotation pipeline generates a robust and reliable catalog of translated ncORFs in mammals.

### ncORFs exhibit signatures of translation and encode putative proteins lacking known domains

We classified ORFs into categories based on community recommendations^2^ and found that the distribution of ORF types closely aligns with previous estimates^18^. In both human and mouse, uORFs, upstream overlapping ORFs (uoORFs), and lncORFs are more prevalent than other types, whereas internal ORFs (iORFs), downstream overlapping ORFs (doORFs), and dORFs are relatively rare (Fig. 2A). Although most ncORFs are short, encoding fewer than 100 amino acids (Fig. 2B), their sequences show clear signatures of translation similar to CDSs. These include preferential usage of AUG start codons (Fig. S3A), enrichment toward 5’ transcript ends (Fig. S3B), and reduced RNA secondary structure^46^ (Fig. S3C) and optimized Kozak contexts^47,48^ around start codons, especially in non-AUG ncORFs (Figs. S3D & S4). Despite these similarities, ncORFs differ markedly from CDSs in both codon and amino acid usage (Figs. S5A-B). This divergence is further supported by their lower tRNA adaptation index values and higher effective number of codons, indicating less optimized codon usage (Figs. S5C-D). Together, these findings show that ncORFs share core sequence and structural features with canonical CDSs, yet maintain distinct codon and amino acid usage profiles.

**Figure 2.**
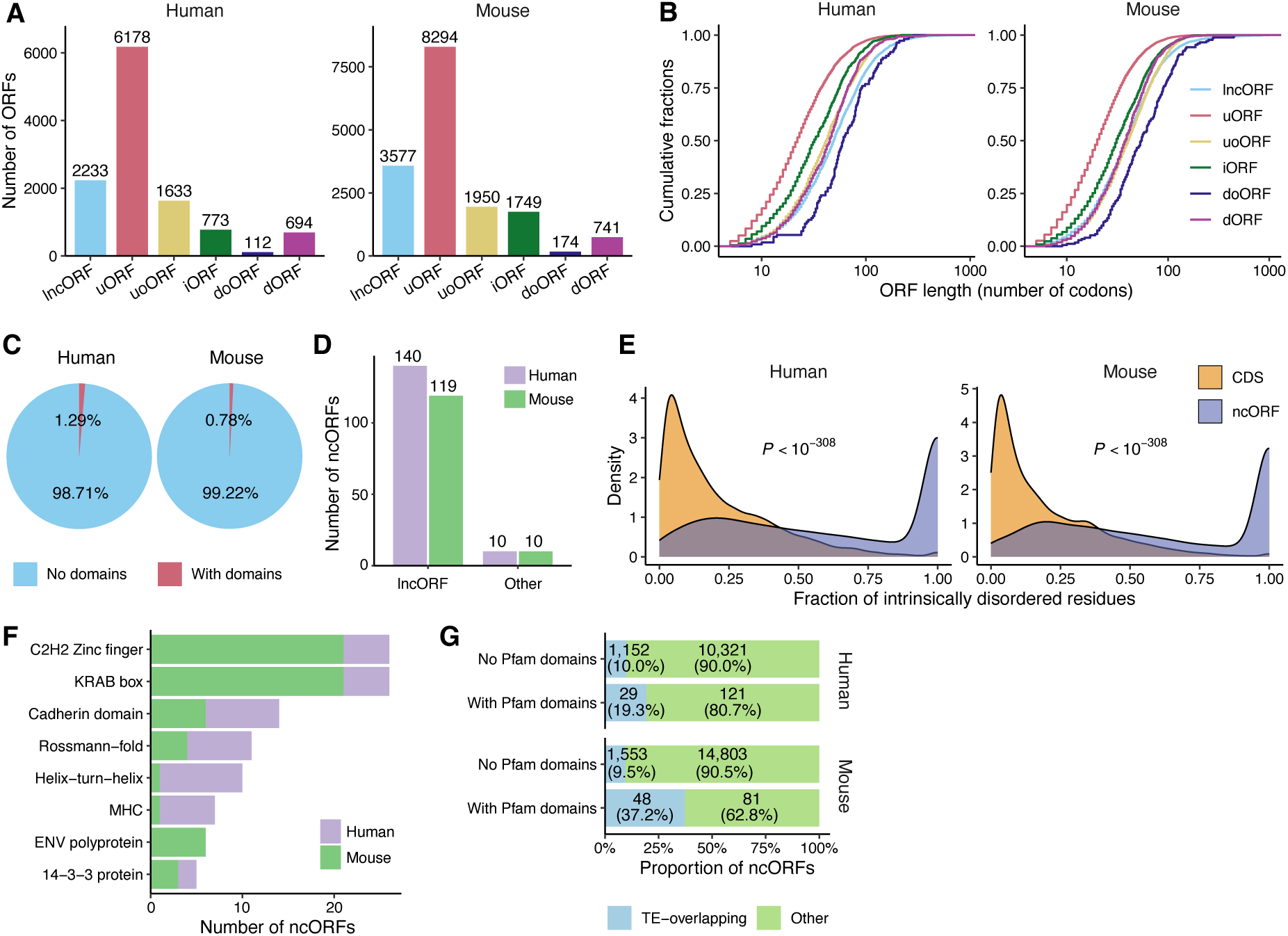
Sequence features of ncORFs and putative ncEPs in humans and mice. (**A**) Number of ncORFs belonging to different categories. (**B**) Distribution of ncORF lengths. (**C**) Proportion of ncEPs that contain known Pfam domains. (**D**) Number of ncORFs with Pfam domains among lncORFs and other ncORF categories. (**E**) Distribution of the proportion of intrinsically disordered residues in CDS- and ncORF-encoded proteins. (**F**) Most frequently identified Pfam domains among human and mouse ncORFs. (**G**) Proportion of ncORFs overlapping with TEs, stratified by presence or absence of Pfam domains. Differences were assessed using Wilcoxon rank-sum tests. CDS, coding sequence; ncORF, non-canonical open reading frame; ncEP, ncORF-encoded protein; TE, transposable element.

Although ncORFs can encode putative proteins, the functional significance of ncEPs remains poorly understood. As an initial step toward functional inference, we systematically searched for annotated protein domains within ncEP sequences, reasoning that the presence of conserved domains would suggest potential biochemical activity. Overall, only 1.29% (150) of human and 0.78% (129) of mouse ncEPs contained at least one known domain (Fig. 2C; Table S5), presumably due to their short lengths relative to canonical proteins. Consistently, domain-containing ncORFs are significantly longer than those without annotated domains (Fig. S6) and are predominantly classified as lncORFs (Fig. 2D). Unlike canonical proteins, which have relatively low intrinsically disordered region (IDR) content (median ± standard error: 14.6±0.28% in humans and 12.7±0.26% in mice), ncEPs exhibit significantly higher IDR proportions (Fig. 2E), with medians of 62.5±1.42% and 60.0±1.27%, respectively. These results indicate that most ncEPs lack structured domains and are instead dominated by IDRs.

Among the eight domains observed in at least five ncORFs, three are DNA-binding domains: zinc finger, Krüppel-associated box, and helix-turn-helix (Fig. 2F). Because transposable elements (TEs) commonly encode DNA-binding proteins required for transposition^49^, we tested whether domain-containing ncORFs preferentially overlap TE-derived sequences. Indeed, domain-containing ncORFs were significantly enriched for TE overlap in both humans and mice (*P* = 5.8×10^−4^ and 3.4×10^−17^, respectively; Fisher’s exact tests; Fig. 2G). Although enrichment alone does not establish origin, it supports a model in which a subset of longer ncORFs may have acquired modular domains through TE-derived sequence contributions.

### Putative ncEPs are selectively constrained

Given the scarcity of known domains in putative ncEPs, a key question is whether they are functional in cells. If at least some of these ncEPs are not merely translational noise or byproducts of translation-dependent functions, we would expect them to exhibit evolutionary signatures of selective constraints similar to canonical CDSs. To test this, we leveraged the recent high-resolution genomic constraint maps derived from both intra- and inter-species comparisons in humans ^50,51^. First, we examined the genomic non-coding constraint of haploinsufficient variation (Gnocchi), which quantifies selective pressure by comparing observed versus expected variants across human genomes. A Gnocchi score above zero indicates purifying selection^50^. Although ncORFs exhibit lower Gnocchi scores than CDSs, their score remain significantly above zero and exceed those of lncRNAs (Fig. S7), suggesting that ncORF sequences are under evolutionary constraint. A hallmark of functional coding regions is codon position-specific constraints, but Gnocchi scores lack the resolution needed to assess such fine-scale features. Therefore, we next analyzed the base-resolution conservation metric PhyloP^52^ calculated from multiple alignments of primate or mammalian genomes^51^. As expected, PhyloP scores in CDSs are substantially higher than those in flanking UTRs and display strong three-nucleotide periodicity, with the third codon position being less conserved than first and second (Fig. 3A). Although ncORFs (excluding those overlapping annotated CDSs) do not show higher overall PhyloP than flanking regions, they also exhibit significant three-nucleotide periodicity in both mammals (Fig. 3A) and primates (Fig. S8), suggesting that at least a subset of ncORFs are subject to similar evolutionary constraints as CDSs.

**Figure 3.**
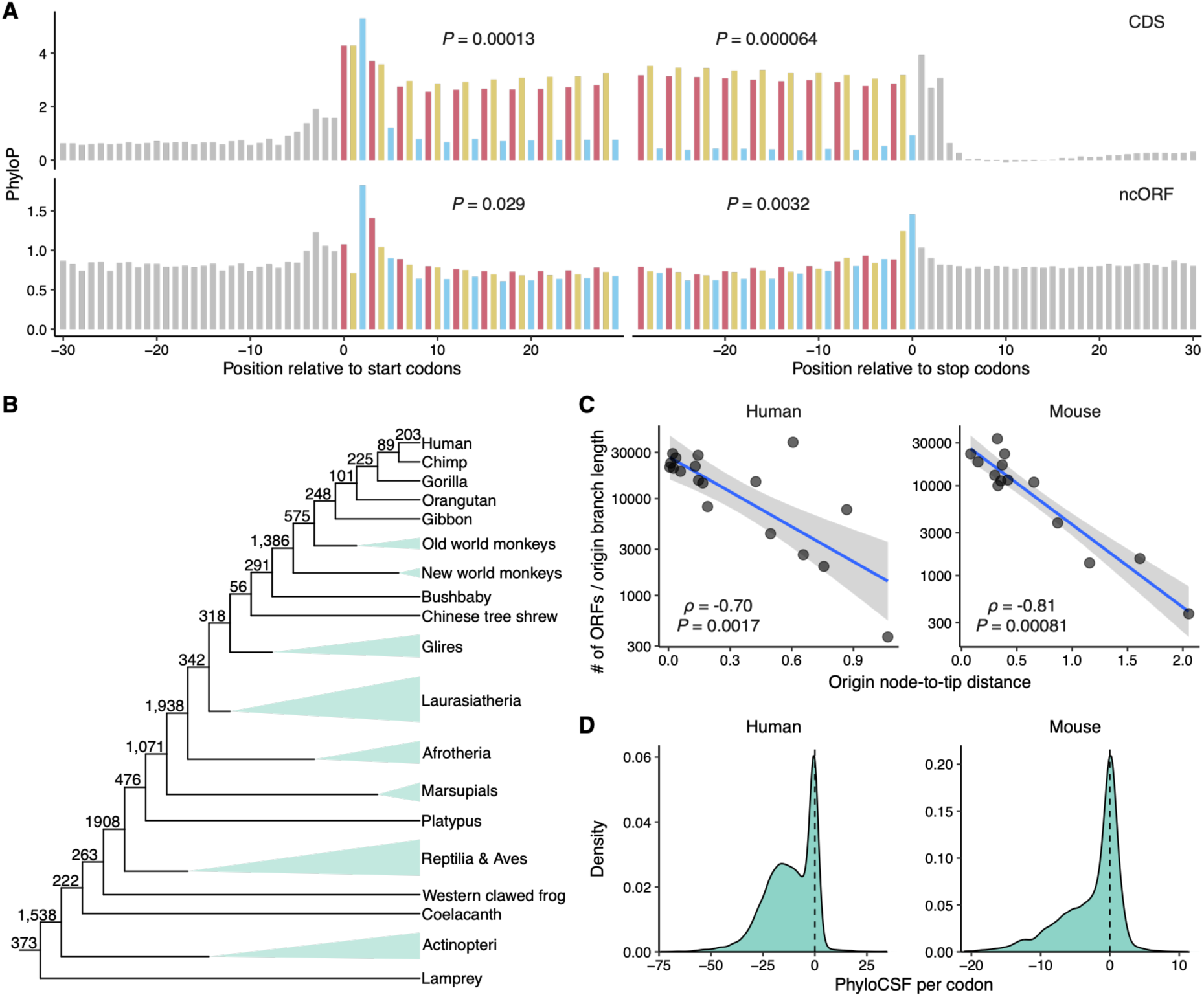
Evolutionary constraints of ncORFs. (**A**) The mean PhyloP scores of 30 base pairs upstream and downstream of the ORF start or stop codons in mammals. Codons were delineated with bars of alternating colors by different frames, and nucleotides in untranslated regions were shown in grey. The significance of three-nucleotide periodicity was assessed by autocorrelation with a lag of three. (**B**) Cladograms of vertebrates with the number of ncORFs originating at each ancestral branch of humans. Triangles indicate species merged into larger clade for visual simplicity. (**C**) Relationship between ORF origination rates and node ages measured as node-to-tip distances. Origination rate was defined as the number of ncORFs that originated on a branch divided by the branch length. Blue lines show linear regression fits, and grey bands represent 95% prediction confidence intervals. Spearman’s correlation is indicated. (**D**) Distribution of ncORF PhyloCSF scores normalized by the number of codons in per ncORF. The dashed line denotes zero. CDS, coding sequence; ncORF, non-canonical open reading frame; ORF, open reading frame.

Motivated by the above results, we next sought to identify individual ncORFs under selective pressure. To this end, we determined their modes of origination^33,34^ and evolutionary history through comparative genomic analyses. Our analysis revealed that most ncORFs are evolutionarily young, with 63.0% (7,319) and 82.8% (13,508) ncORFs in humans and mice originating within the mammalian lineage, respectively (Figs. 3B & S9). In line with previous studies^33,34^, we estimate that at least 51.7% of human and 50.8% of mouse ncORFs arise *de novo* (Table S6). We also observed that the number of ncORFs originating at each ancestral node, when normalized by branch length, decreases with increasing node-to-tip distance (Fig. 3C), reflecting dynamic turnover and reduced long-term retention of ncORFs. To assess whether individual ncORFs are constrained for coding, we analyzed codon substitution patterns within ncORFs since origination^33^ with PhyloCSF^53^. PhyloCSF estimates the log-likelihood ratio of a given alignment under coding versus non-coding models; positive values indicate stronger similarity to conserved coding sequences. We found 13.8% (1,608) of human and 32.0% (5,281) of mouse ncORFs have positive PhyloCSF scores (Fig. 3D), and 10.9% (1,264) of human and 22.6% (3,721) of mouse ncORFs even have PhyloCSF exceeding 10 (Table S6), providing strong evidence that these sequences may encode functional ncEPs.

Evolutionary dynamics can also provide insights into the function of ncORFs, as functionally constrained ncORFs are less likely to be lost over time. To quantify the conservation of each ncORF since its origination, we developed a local branch length score (BLS) (Fig. 4A), adapting the “global” BLS metric previously used to assess uORF conservation across full phylogenies^48,54^. Unlike global BLS, local BLS is designed to detect lineage-specific constraint, making it better suited for identifying functional ncORFs that may be restricted to certain evolutionary branches. Using a local BLS threshold of 0.9, we identified 16.3% (1,889) of human and 24.8% (4,087) of mouse ncORFs as exhibiting lineage-specific conservation (Figs. 4B & S9). Notably, these ncORFs are significantly more likely to show positive PhyloCSF scores (Fig. 4C), reflecting the effectiveness of local BLS. In total, we identified 681 human and 1,622 mouse ncORFs that are both lineage-specifically conserved and show evidence of coding potential, making them strong candidates for future functional validation. Together, these findings suggest that a subset of ncORFs may encode functional proteins rather than translational byproducts.

**Figure 4.**
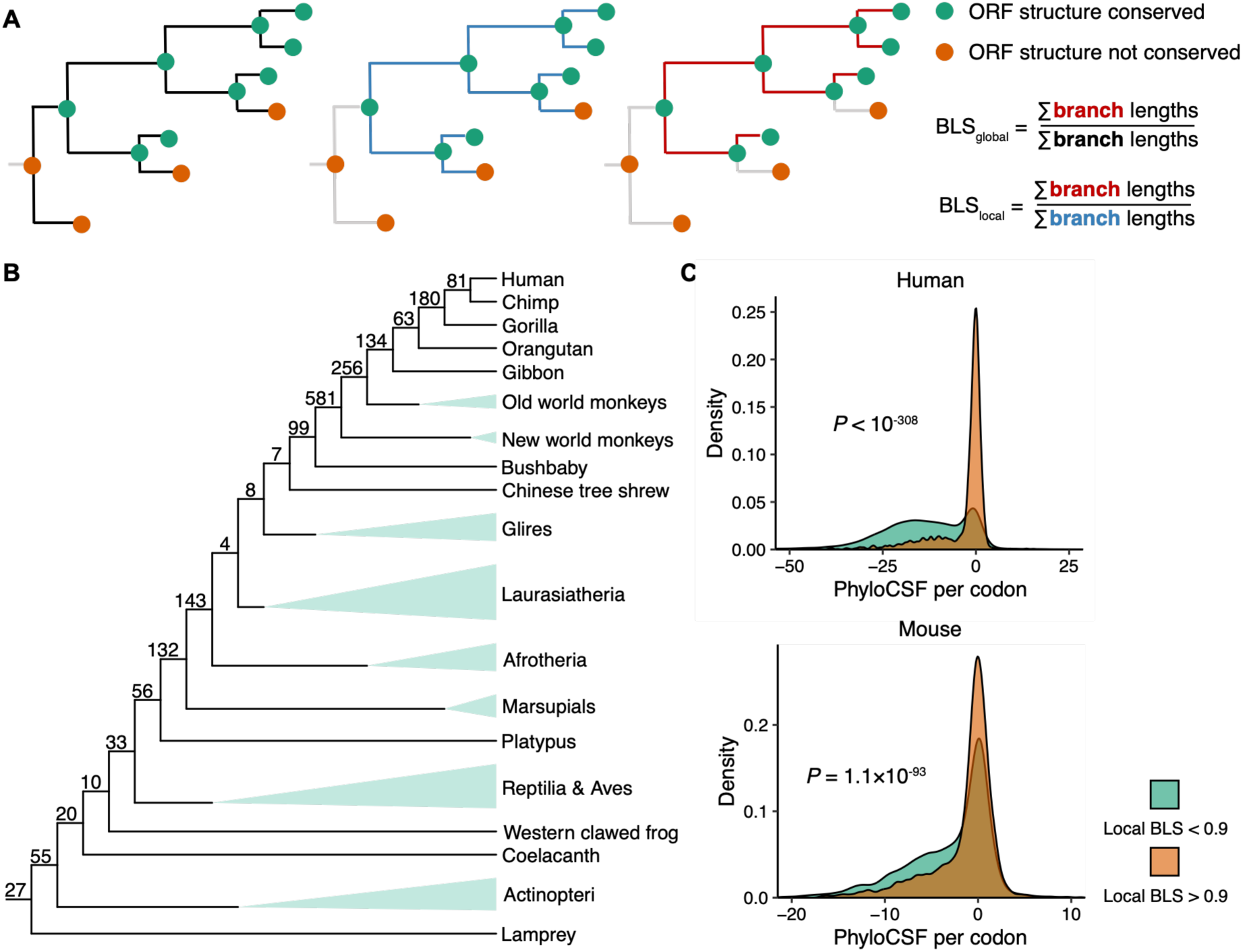
Lineage-specifically conserved ncORFs. (**A**) Schematic illustration for BLS calculation. (**B**) Cladograms of vertebrates showing the number of lineage-specifically conserved ncORFs (local BLS > 0.9) at each ancestral node for humans and mice. (**C**) Distribution of PhyloCSF scores per codon for lineage-specifically conserved ncORFs compared to all other ncORFs. BLS, branch length score; ncORF, non-canonical open reading frame; ORF, open reading frame.

### Expression of ncORFs is shaped by their evolutionary dynamics

Gene expression is a tightly regulated process, and non-functional products are unlikely to be robustly expressed. To assess the expression of ncORFs, we examined their translation across diverse tissues and cell lines (Table S7). On average, 71.9% of human and 48.5% of mouse ncORFs are translated with reads per kilobase per million mapped reads (RPKM) ≥ 1, which is comparable to or slightly exceeds the proportions observed for CDSs (Fig. S10). To investigate the influence of ncORF evolution on their expression, we analyzed mean translation levels of ncORFs across different samples. We found that ncORFs with higher local BLS tend to be more highly translated in both species (Fig. S11A), whereas coding potential showed only a minor influence (Fig. S11B). To disentangle the effects of evolutionary constraint and gene age, we stratified ncORFs by origination nodes and examined their translation levels in relation to local BLS and PhyloCSF. Although older ncORFs are generally more highly translated, those with higher local BLS scores consistently show elevated translation levels compared to their age-matched counterparts (Figs. 5A & S12A). This suggests that lineage-specific conservation is associated with enhanced and possibly optimized translation, potentially reflecting acquired functional roles. Conversely, the relationship between coding potential and translation is more nuanced: younger ncORFs with positive PhyloCSF scores tend to be more highly translated, while the opposite trend is observed in older ncORFs (Figs. 5B & S12B). These patterns point to complex interplays between evolutionary constraint, coding potential, and translational output, likely reflecting divergent selective pressures across different ncORF age groups.

**Figure 5.**
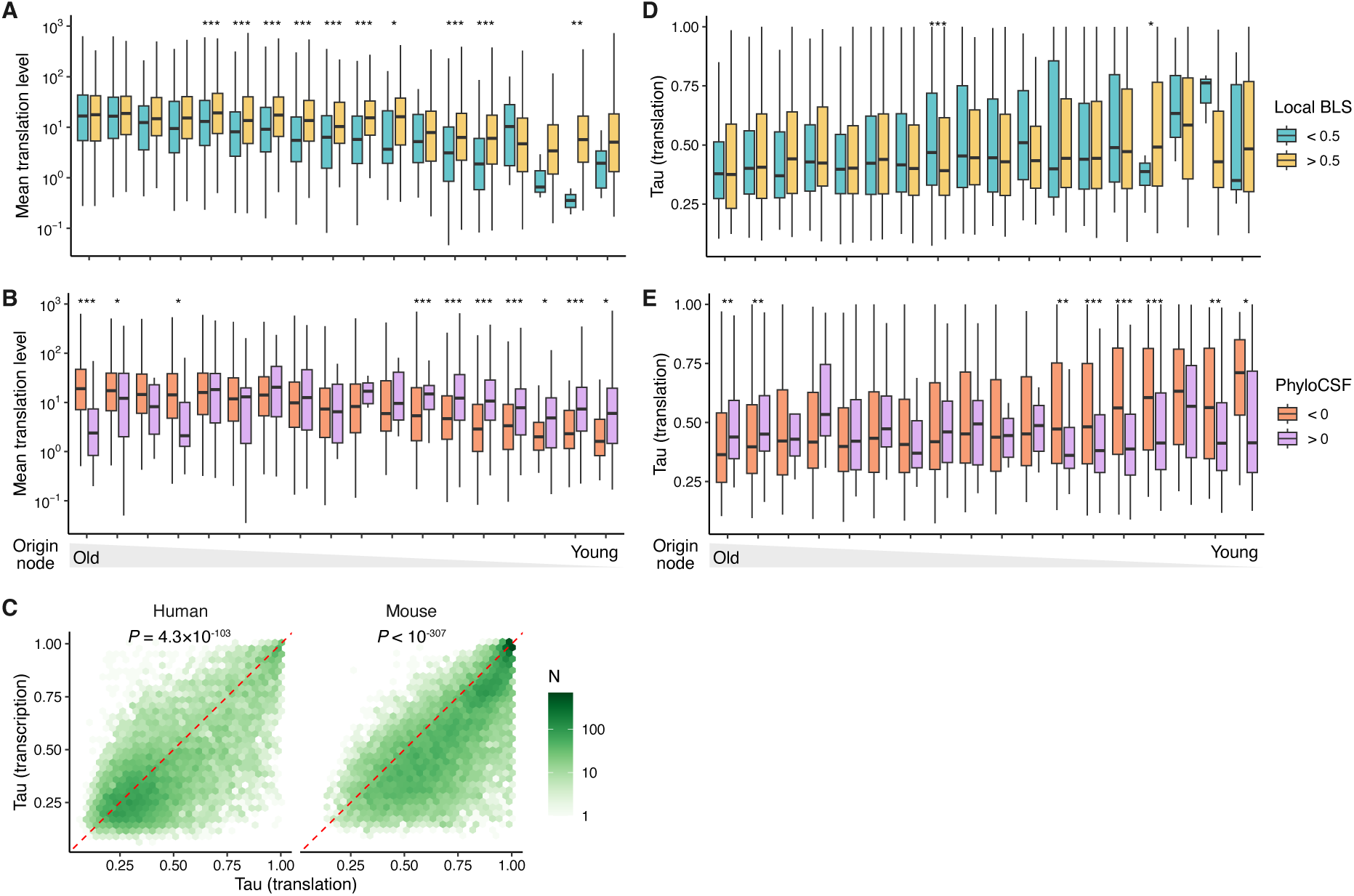
Evolutionary dynamics of ncORF expression. (**A**) Distribution of mean translation levels of human ncORFs grouped by their origin nodes and further stratified by local BLS. Statistical significance was assessed using Wilcoxon rank-sum tests. ***, *P* < 0.001; **, *P* < 0.01; *, *P* < 0.05. (**B**) Similar to (A) but ncORFs are further stratified by local BLS. (**C**) Relationship between ncORF tissue specificity at translation and transcription levels. Differences were determined with Wilcoxon signed-rank tests. (**D-E**) Similar to (A) and (B) but showing the distribution of tissue specificity at the translation level. CDS, coding sequence; ncORF, non-canonical open reading frame; ORF, open reading frame.

While the above analyses revealed the overall trends in ncORF translation, they overlooked the heterogeneity across tissues and cell types. We found that ncORF translation exhibits strong tissue specificity (Fig. S13), which may stem from either transcriptional bias in ncORF-containing genes or post-transcriptional regulation specific to ncORFs. To disentangle these factors, we examined RNA levels of ncORF-containing genes across different tissues^55,56^. Although transcriptional enrichment in testis was evident in both humans and mice (Fig. S14), testis-biased translation was observed only in mice (Fig. S13). Moreover, ncORF translation displayed significantly higher tissue specificity than the transcription of their host genes in both species (Fig. 5C), indicating a substantial post-transcriptional component to their regulation. We next assessed how evolutionary characteristics shape tissue specificity. Older ncORFs tend to have lower tissue specificity than younger ones, suggesting they are more broadly translated (Figs. 5D & S12C). Intriguingly, coding potential exhibits opposite effects depending on evolutionary age: younger ncORFs with higher coding potential are more broadly expressed, whereas older ones show increased tissue specificity (Figs. 5E & S12D). These patterns may help to explain the lower average translation levels observed for older ncORFs across tissues. A similar trend was also evident for ncORFs with high local constraint scores in mice (Fig. S12C). Collectively, these results support a model in which ncORFs initially evolve widespread expression to establish interactions and avoid loss^57^, but may later become functionally specialized, resulting in more restricted expression.

### Extensive co-translation between ncORFs and canonical CDSs

Given our earlier observation that many ncEPs lack modular domains and may act via interactions with canonical proteins, we next explored the landscape of co-translation between ncORFs and CDSs across different tissues or cell types as we expect that interacting proteins tend to be translated simultaneously spatiotemporally. Using weighted gene co-expression network analysis (WGCNA)^58^, we identified 61,513 human and 55,449 mouse ncORF-CDS co-translated pairs. As expected, ncORFs in mRNAs tend to co-translate with the main CDS of the same gene, even after excluding CDS-overlapping ones (Fig. S15), consistent with previous experimental findings^28^. Among non-overlapping co-translating ncORFs, only 10 in humans were dORFs; all others in both species were uORFs. Of note, co-translation does not contradict the canonical model in which uORFs repress downstream CDS translation. Co-translation reflects concurrent ribosome occupancy driven by overall initiation rates at the 5’ end, whereas repression concerns the fraction of initiating ribosomes that reach the CDS. Thus, absolute translation of both ORFs may increase without altering the relative inhibitory effect of the uORF^59,60^.

ncORF-CDS co-translation forms a highly modular bipartite network, in which one CDS may be linked to multiple ncORFs and vice versa (Fig. 6A-B). Network analysis revealed that hub CDS nodes—those with the most connections—in the largest modules are significantly enriched for functional gene categories (Fig. 6C-D). For instance, the largest module in humans is associated with chromatin-related proteins, while the second largest module in humans and the largest one in mice are linked to extracellular matrix organization. Notably, ancient ncORFs (those originating before the last common ancestor of mammals) are significantly more likely to co-translate with CDSs than younger ones (Fig. 6E), potentially reflecting a greater likelihood of acquiring functional interactions over evolutionary time. In summary, our analyses revealed that evolutionary dynamics of ncORFs have widespread impacts on their translation and potential function.

**Figure 6.**
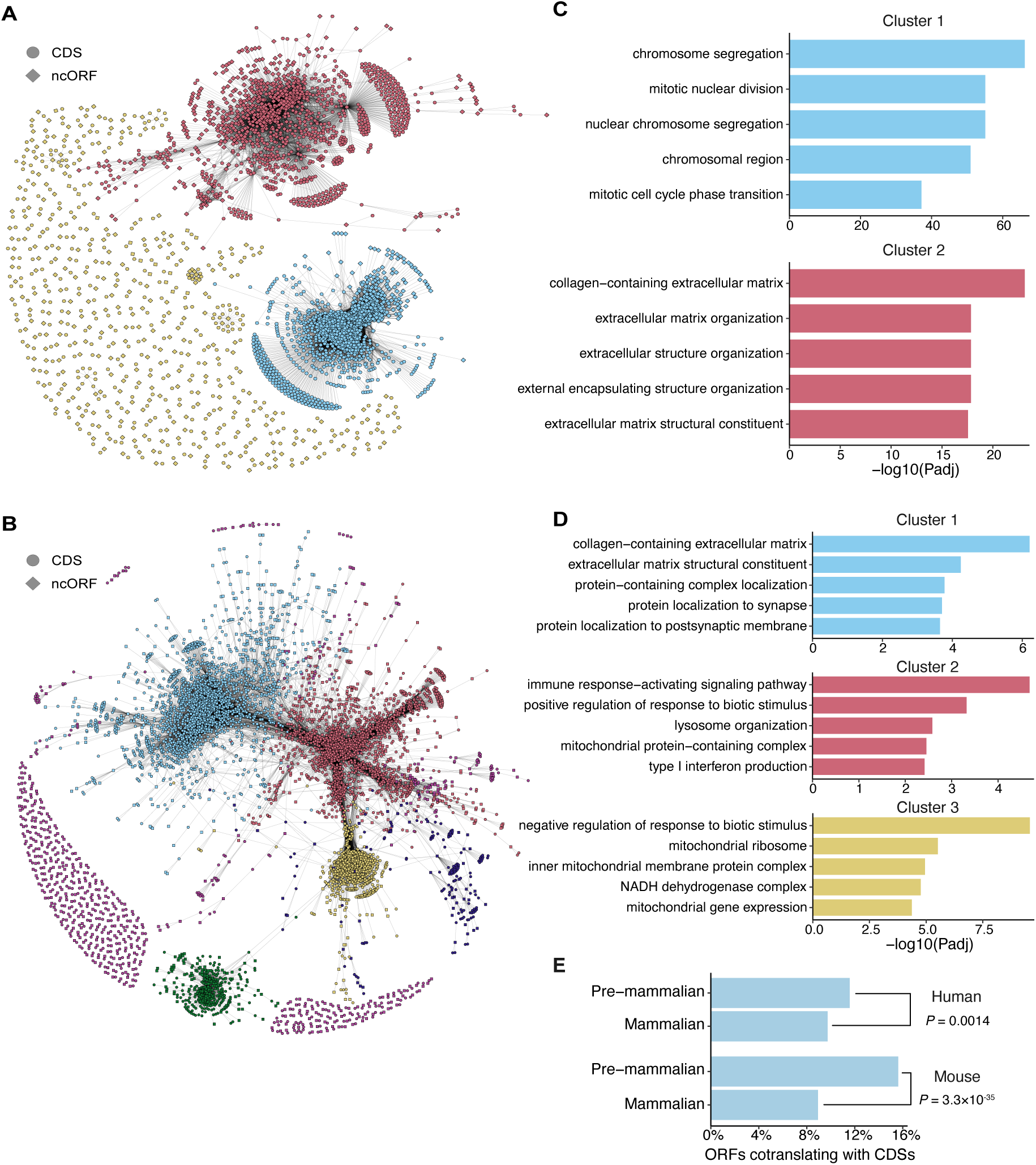
ncORF-CDS co-translation network. (**A**) Bipartite network of ncORF-CDS co-translation in humans. (**B**) Bipartite network of ncORF-CDS co-translation in mice. ncORFs and CDSs are represented by different shapes and colored according to their cluster membership. Only the largest clusters were highlighted (top two in humans and top five in mice). (**C**) Top five enriched gene ontology terms for each of the two largest clusters in the human network shown in (A). (**D**) Top five enriched gene ontology terms for each of the three largest clusters in the mouse network shown in (B). (**E**) Proportion of ncORFs cotranslating with CDSs among ancient (pre-mammalian origin) versus younger (mammalian-specific) ncORFs. Differences were tested with Fisher’s exact tests. CDS, coding sequence; ncORF, non-canonical open reading frame; ORF, open reading frame.

## Discussion

Reliable annotation of translated ncORFs is critical for understanding their functional roles and evolutionary significance. In this study, we developed a standardized pipeline for the annotation of translated ncORFs and applied it to both humans and mice due to the extensive publicly available Ribo-Seq datasets. Compared to previous efforts, our annotation framework offers several key improvements. First, we selected only high-quality libraries with sufficient read coverage and acceptable in-frame RPF rates. Our earlier work demonstrated that both library size and in-frame rates impact the reproducibility and accuracy of ncORF predictions^20^. This step helps mitigate false annotations arising from suboptimal data quality. Second, although many previously characterized functional ncEPs are associated with cancer-related ncORFs^1^, ncORFs may also have important functions under physiological conditions. Importantly, such ncORFs are more likely to be subject to evolutionary constraints. Therefore, we restricted our analysis to normal cells and tissues, allowing us to identify more biologically relevant and conserved ncORFs. Third, given the variable performance across ncORF detection methods and the low reproducibility of predictions between biological replicates^20^, we implemented stringent reproducibility criteria. Specifically, we retained only ncORFs detected by multiple methods and across independent datasets, enhancing the robustness of our final annotation. Finally, we applied a suite of well-validated filtering metrics from prior studies^39–41^ to minimize the inclusion of false positives, including those arising from translational noise^61,62^ or non-ribosomal contamination. As a result, our annotated ncORFs show significant enrichment for GENCODE ncORFs, especially those with MS evidence or reproducible detection across independent studies. These features collectively underscore the reliability and utility of our annotation for future functional and evolutionary studies of non-canonical translation.

However, our catalog may remain incomplete due to the uneven distribution of publicly available Ribo-seq datasets across tissues and cell types. Broader and more balanced tissue coverage will be necessary to generate a more comprehensive and less biased ncORF annotation, which would also enhance the cotranslation analysis. Additionally, our proteomic validation relies on previously compiled sets of MS-supported ORFs rather than de novo reanalysis of raw mass spectrometry datasets. Incorporation of candidate ncORFs into reference proteome databases prior to large-scale, standardized searches of public MS runs would substantially improve detection sensitivity and provide stronger evidence for their translational products^45^.

From the perspective of new gene evolution, ncORFs represent a transitional stage between non-coding DNA/RNA and protein-coding genes^63,64^. A major challenge in reclassifying ncORFs as bona fide coding genes lies in determining whether the putative proteins they encode are functionally relevant. Our analysis of ncEPs revealed that most lack recognizable protein domains and are enriched for IDRs. Therefore, we propose that many ncORFs may function in a non-autonomous manner by interacting with other proteins to exert their effects. This hypothesis aligns with prior studies showing physical interactions between ncEPs and known proteins^28,29,34^. Supporting this view, our WGCNA analysis uncovered widespread co-translation between ncORFs and canonical CDSs, suggesting potential interaction partners and functional contexts.

We also assessed whether ncORFs are subject to evolutionary constraints consistent with functional protein coding. Using both population-level and cross-species analyses, we found that a substantial fraction of ncORFs bear signatures of purifying selection, in line with previous studies^33,59,60,65,66^. Notably, their conservation profiles exhibit nucleotide-level features characteristic of canonical CDSs, supporting the interpretation that selection acts on the encoded peptide products rather than merely on translational activity per se. Furthermore, we reconstructed the evolutionary trajectories of thousands of ncORFs in both humans and mice and observed sustained coding potential since their origination. Complementing these findings, we identified a subset of ncORFs that preserve intact ORF structures across most descendant species, suggesting ongoing purifying selection against disabling mutations. These lineage-specifically conserved and potentially coding ncORFs likely contribute to clade-specific functional innovation and represent compelling candidates for future functional characterization.

We further explored how evolutionary history shapes ncORF translation across tissues or cell types. Our results indicate that older ncORFs are more broadly and highly translated and have established more extensive co-expression with canonical CDSs. Moreover, when comparing ncORFs of similar age, those that are lineage-specifically constrained consistently exhibit higher translation levels, reinforcing their likely functional importance. Interestingly, we found that younger ncORFs with strong evolutionary constraints tend to have lower tissue specificity in their translation, potentially reflecting a phase of widespread expression that facilitates functional recruitment. In contrast, older ncORFs with positive coding potential tend to display increased tissue specificity, implying functional specialization over evolutionary time. Collectively, these findings support a model in which ncORFs evolve from broadly expressed, weakly functional elements into specialized, conserved effectors—though further empirical work is needed to validate this hypothesis.

These patterns should nevertheless be interpreted in light of the heterogeneous nature of the public Ribo-seq datasets used in our analysis. The datasets differ in experimental protocols, sequencing depth, read quality, tissue and cell-type coverage, and other technical factors that can influence the sensitivity and specificity of ncORF detection and quantification. In particular, ncORFs with low or highly restricted translation may be preferentially missed in datasets with limited depth or suboptimal quality, potentially affecting estimates of translation breadth and tissue specificity. Systematic and standardized Ribo-seq profiling across matched tissues, developmental stages, and species will be required to more fully resolve the evolutionary dynamics of ncORF translation.

In conclusion, we provide a comprehensive annotation of translated ncORFs in humans and mice, offering a valuable resource for functional and evolutionary genomics. Our integrated analysis of sequence evolution and translational landscape sheds light on the evolutionary trajectory and functional integration of ncORFs. The consistency of these patterns in both species suggests that our findings may generalize to other mammals. However, realizing the full biological significance of ncORFs, particularly lineage-specific ones, will require experimental validation. Recent advances in high-throughput screening technologies^19,29,32,67^ offer a promising path forward. Lastly, although our study focused on humans and mice due to the availability of high-quality Ribo-Seq data, we anticipate that our annotation will serve as a training resource for machine learning models aimed at predicting translated ncORFs in non-model organisms, thus broadening our understanding of genome coding potential across species.

## Methods

### Quality control and preprocessing of Ribo-Seq libraries

Ribo-Seq libraries from normal human or mouse samples without genetic modifications or perturbations (Table S7) were downloaded from the European Nucleotide Archive (ENA) or the Short Read Archive (SRA). For ncORF annotation, libraries containing fewer than five million reads were excluded. Libraries derived from cancer tissues or cell lines were also removed based on sample metadata or the associated publications. Next, adapter sequences and any random nucleotide additions were trimmed using Cutadapt (v3.7)^68^ and reads shorter than 18 nucleotides were discarded. Remaining reads were aligned to the human or mouse reference genome (GRCh38 or GRCm39, ENSEMBL^69^ release 107) using the STAR aligner (v2.7.10a)^70^. To assess library quality, the proportion of in-frame ribosome-protected fragment (RPF) reads was calculated using the PSite package^71^. Libraries with ≥60% in-frame RPFs were classified as high quality. Only samples containing at least two high-quality libraries were retained for ncORF annotation (Table S1). For the expression analyses, all the downloaded Ribo-Seq libraries were used to cover more samples.

### ncORF annotation

For each Ribo-Seq library, translated ORFs initiating with AUG, CUG, GUG, or UUG and encoding at least five amino acids were predicted using PRICE^36^, RiboCode^37^, and Ribo-TISH^38^, as described previously^20^. To define novel ORFs, annotated CDSs and their truncation or extension variants (i.e., ORFs with the same start or stop position as annotated CDSs) were excluded. To eliminate likely false positives resulting from non-ribosomal contamination or stochastic molecular errors, we calculated RRS^40,41^ and FLOSS^39^ of each ncORF. RRS measures the drop in RPF coverage after stop codon—a key signature of active translation beyond three-nucleotide periodicity^40,41^, whereas FLOSS assesses whether the RPF length distribution resembles that of canonical CDSs, helping to distinguish authentic ORFs from non-ribosomal signals^39^. In each library, uniquely mapped RPFs were assigned to the corresponding P-site, and genome-wide P-site coverage was computed with the PSite package. Similar to previous studies^40,41^, RRS was calculated as the sum of coverage across the five codons preceding the stop codon divided by the sum across the five trinucleotides following the stop codon. FLOSS was calculated by first deriving a reference RPF length distribution vector (*P_ref_*) using all RPFs uniquely mapped to annotated CDSs. For each ORF (ncORF or CDS), a read length frequency vector (*P_ORF_*) was computed, and FLOSS was calculated as ∑|*P_ref_* − *P_ORF_*|/2. The empirical upper bound for acceptable FLOSS was set using Tukey’s method (i.e., the third quartile + 1.5 × inter-quartile range of FLOSS values across all CDSs)^39^. ncORFs with fewer than 10 P-site-mapped RPFs, RRS ≤ 1, or FLOSS values exceeding the empirical upper bound were filtered out.

Integration of ncORF prediction across different libraries of the same species was nontrivial due to extensive overlap among ncORFs in genomic coordinates. To address this issue, we used a graph-based clustering approach. We created a graph where two ORF vertices were connected if they share the same start or stop codon. Then, each set of connected ncORFs in the graph represents a cluster of overlapping ncORFs. To ensure reproducibility, only clusters identified by ≥2 methods in ≥2 libraries were retained, yielding 13,071 clusters in humans and 17,674 clusters in mice. To ensure compatibility with the latest reference gene annotations (ENSEMBL release 111), we removed any ncORFs that were newly classified as annotated CDSs. A manual inspection of RPF signals using the GWIPS-viz Ribo-Seq track^72^ in the UCSC Genome Browser^73^ revealed additional spurious ncORFs overlapping in-frame with CDSs from other transcripts. These were likely misidentified due to shared translation signals and were conservatively excluded using custom scripts, leaving 11,623 clusters in humans and 16,485 in mice.

To facilitate downstream analyses, a representative ncORF was selected for each cluster using a two-step strategy. First, a representative transcript was selected by prioritizing those with matched annotation from NCBI and EMBL-EBI (MANE)^42^ (for humans) or APPRIS principal^74^; otherwise, the longest transcript was chosen. Then, among ncORFs within the representative transcript, a single ncORF was selected based on start codon hierarchy (AUG > CUG/GUG > UUG) following empirical evidence^28,36^. If multiple ncORFs shared the same preferred start codon, the longer one was chosen. Finally, we classified ncORFs according to their positions relative to annotated transcript features. ncORFs located in lncRNAs were designated as lncORFs. Among ncORFs in protein-coding transcripts, those located entirely within the 5′ UTR were classified as uORFs, whereas those initiating in the 5′ UTR and extending into the CDS were classified as uoORFs. ncORFs with both start and stop codons within the annotated CDS were classified as iORFs, whereas those initiating within the CDS and terminating in the 3′ UTR were classified as doORFs. Those initiating in the 3′ UTR were classified as dORFs.

For validation, our ncORFs were compared with previously annotated GENCODE ncORFs^2^ and those with MS evidence compiled previously^20^. Tier annotations of GENCODE ncORFs were obtained from a recent study^45^. Known functional ncORFs and their genomic coordinates were manually curated from published literature (Table S4). To account for possible proteoforms arising from alternative initiation and splicing, we considered an ncORF to be recovered if either its start or stop codon was identical to any ncORF in our annotation.

### ncORF sequence analyses

The secondary structure surrounding each ORF (ncORF or CDS) start codon was predicted with RNAfold^75^ on a 150 nt window extracted from the transcript in which the ncORF resides starting from the 30^th^ nucleotide upstream of the start codon. For each position within this window, the overall pairing probability was calculated by dividing the number of ORFs with a base-paired residue at that position by the total number of ORFs analyzed. The tRNA adaptation index and effective number of codons for ncORFs and CDSs were calculated using the “cubar” R package^76^. tRNA gene annotations in humans and mice were obtained from GtRNAdb (release 22)^77^.

### Putative ncEPs analyses

Known domains within ncEPs were identified using InterProScan (v5.64-96.0)^78^ with default parameters. Repetitive sequence annotations were downloaded from the UCSC Table Browser. BEDTools (v2.31.1)^79^ was used to identify TE-overlapping ncORFs. IDRs were predicted with IUPred3^80^ with analysis type set to “short disorders” and smoothing disabled. For each ORF, residues with disorder tendency score > 0.5 were considered disordered, and the IDR fraction was calculated as the proportion of disordered residues relative to the ORF length.

### Gnocchi Score and PhyloP of ncORFs and CDSs

Genome-wide Gnocchi scores computed using 1 kb sliding windows (with 100bp step size) were downloaded from the gnomAD database^50^. For each base pair, the Gnocchi score was defined as the average across the ten overlapping 1kb windows that covered that position using BEDTools. The final Gnocchi score for each ORF (ncORF or CDS) was obtained by averaging the scores of all base pairs within the ORF.

Base-wise PhyloP scores computed across 441 mammals or 239 primates^51^ were downloaded from the UCSC Genome Browser. For each ncORF or CDS, PhyloP values were extracted for a ±30 bp window surrounding the start and stop codons using custom scripts. These values were then averaged over different ORFs to generate metagene profiles for visualization. To evaluate three-nucleotide periodicity, the metagene profiles were subjected to autocorrelation analysis using the “ac.test” function from the “testcorr” R package^81^. Statistical significance was assessed using the robust *t̃* statistics.

### ncORF origination node and mode

Whole genome alignments and corresponding species trees were downloaded from the UCSC Genome Browser (GRCh38 100-way for humans and GRCm39 35-way for mice). The origination node and mode for each ncORF were inferred following previous approaches^33,34^. For each ORF, the sequence from the reference genome and its orthologous regions in other species were extracted. Orthologous sequences covering less than 50% of the ncORF length relative to the reference species were excluded. The remaining sequences were realigned with MAFFT (v7.520)^82^ with default parameters. The corresponding gene tree was derived by pruning the species tree using Gotree (v0.4.3)^83^ with default settings. Ancestral sequence reconstruction was performed using FastML (v3.11)^84^ with parameters “--seqType nuc --SubMatrix HKY --OptimizeBL no --indelReconstruction ML”. An ORF structure was considered intact in an orthologous or ancestral sequence if one of the first three codons was a start codon and at least 70% of the 5’ region of the reference ncORF was free of in-frame stop codons. The origination node was defined as the most ancient ancestral node that retained an intact ORF structure. An ncORF was classified as *de novo* if the origination node had an ancestor covering at least 50% of the ncORF sequence but lacking an intact ncORF structure. Otherwise, the origination mode was classified as uncertain. The distance from an ancestral node to the reference species (tip node) was calculated by summing branch lengths along the connecting path using the ape R package (v5.8)^85^.

### ncORF BLS and PhyloCSF

For each ncORF, global BLS was calculated as the ratio of the total branch length of the subtree connecting all leaves with intact ORF structures to the total branch length of the full species tree as in our previous study^48^ with custom scripts. Local BLS was calculated similarly, but the denominator was replaced by the total branch length of the subtree defined by the origination node and all of its descendant nodes (including leaves). To evaluate coding potential, the alignment of all species descending from the origination node was converted into Multiple Alignment Format after removing the stop codon. PhyloCSF scores were then computed using the “score-msa” module of PhyloCSF++^86^.

### ncORF expression analysis

To quantify translation levels of ncORFs and CDSs in Ribo-Seq libraries, we assigned uniquely mapped reads to corresponding P-site positions with the PSite package. The read count for each ORF was calculated as the total number of RPFs whose P-sites fell within the ORF. To reduce bias caused by RPF enrichment at ORF termini, reads mapping to start or stop codons were excluded. For CDSs, only the count of the representative transcript of each gene as defined above was used for downstream analyses. For samples with multiple libraries, read counts were aggregated prior to normalization. Sample-specific size factors were estimated using CDS counts and applied to normalize both ncORF and CDS counts across samples using DESeq2^87^. RPKM for an ORF was calculated as normalized read counts / ORF length (nt) / library size × 10^9^, where library size was the median of total read counts across samples to ensure cross-sample comparability. An ORF (ncORF or CDS) was considered translated if its RPKM was ≥ 1. To evaluate tissue specificity of ncORF translation, the normalized RPF count *X_k_* of an ORF in sample *k* was log-transformed as *x_k_* = *log*(*X_k_* + 1). Then, tissue specificity index^88^ was calculated as 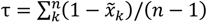, where 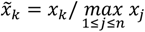 and *n* is the total number of samples. To quantify RNA levels, raw reads of deep transcriptome profiling across different human or mouse tissues from previous studies^55,56^ were downloaded from SRA. Gene expression levels (transcript per million, TPM) were estimated using salmon (v1.10.3)^89^ with parameters “--seqBias --gcBias --posBias -l A”.

WGCNA analysis^58^ of ncORF and CDS translation was performed using the PyWGCNA package (v2.0.5)^90^ with default settings. Only ORFs with normalized RPF counts ≥ 10 in more than half of the samples were included. Pairwise cotranslation relationships were quantified using the topological overlap measure (TOM) derived from the resulting networks. Because the distributions of TOM values differed between the human and mouse networks following species-specific soft-threshold selection, empirical TOM cutoffs were determined separately for each species. Two ORFs were considered cotranslated if their TOM exceeded 0.1 (human) or 0.01 (mouse).^58^ To test whether co-translation is enriched between ncORFs and CDSs from the same transcript, gene identities of CDSs were randomly permuted among all co-translated ncORF–CDS pairs. This procedure was repeated 1,000 times, and the empirical *P* value was calculated as (*m* + 1) / 1,000, where *m* is the number of permutations in which the number of same-gene pairs was greater than or equal to the observed count. The bipartite network of ncORF-CDS co-translation was clustered using the Leiden algorithm^91^ implemented in igraph (v2.1.3)^92^ by optimizing modularity with a resolution of 0.2. The top 20% of nodes by betweenness centrality were designated as hub nodes. Gene ontology analysis of hub genes in the largest modules was conducted using ClusterProfiler (v4.12)^93^.

## Code availability

All analyses were primarily conducted using R software (v4.3). Unless otherwise stated, statistical tests are two-sided. The code and ncORF catalog are available at https://github.com/gxelab/ncorf_mammals and custom scripts used for ncORF classification and evolutionary analyses can be found at https://github.com/gxelab/scripts.

## Data availability

Ribo-Seq libraries for ncORF annotation were obtained from the ENA or SRA, with accession codes listed in Table S1. Libraries used for ncORF and CDS translation quantification were listed in Table S7. RNA-Seq libraries for gene expression profiling across different human or mouse tissues were from previous studies^55,56^ and are available with ENA accession numbers E-MTAB-2836 and E-MTAB-10276, respectively. Whole genome alignment across vertebrates, annotation of genomic repetitive elements, and base-wise PhyloP in primates and mammals^51^ were downloaded from the UCSC Genome Browser^73^. Gnocchi scores were downloaded from gnomAD^50^. All the other data were presented in the manuscript or supplementary tables and are available from the corresponding author upon reasonable request.

## Acknowledgments

We thank Dr. Chuan Yan and all the Zhang lab members for helpful discussion. This study was supported by the National Natural Science Foundation of China (32200433) and the Fundamental Research Funds for the Central Universities (LZUJBKY-2022-2). We thank the Supercomputing Center of Lanzhou University for providing computational resources.

## References

1. Wright, B.W., Yi, Z., Weissman, J.S., and Chen, J. (2021). The dark proteome: translation from noncanonical open reading frames. Trends in Cell Biology. 10.1016/j.tcb.2021.10.010.

2. Mudge, J.M., Ruiz-Orera, J., Prensner, J.R., Brunet, M.A., Calvet, F., Jungreis, I., Gonzalez, J.M., Magrane, M., Martinez, T.F., Schulz, J.F., et al. (2022). Standardized annotation of translated open reading frames. Nature Biotechnology 40, 994–999. 10.1038/s41587-022-01369-0.

3. Azam, S., Yang, F., and Wu, X. (2025). Finding functional microproteins. Trends Genet 41, 107–118. 10.1016/j.tig.2024.12.001.

4. Baena-Angulo, C., Platero, A.I., and Couso, J.P. (2025). Cis to trans: small ORF functions emerging through evolution. Trends Genet 41, 119–131. 10.1016/j.tig.2024.10.012.

5. Hofman, D.A., Prensner, J.R., and van Heesch, S. (2025). Microproteins in cancer: identification, biological functions, and clinical implications. Trends Genet 41, 146–161. 10.1016/j.tig.2024.09.002.

6. Orr, M.W., Mao, Y., Storz, G., and Qian, S.B. (2020). Alternative ORFs and small ORFs: shedding light on the dark proteome. Nucleic Acids Research 48, 1029–1042. 10.1093/nar/gkz734.

7. Lu, S., Zhang, J., Lian, X., Sun, L., Meng, K., Chen, Y., Sun, Z., Yin, X., Li, Y., Zhao, J., et al. (2019). A hidden human proteome encoded by ‘non-coding’ genes. Nucleic Acids Res 47, 8111–8125. 10.1093/nar/gkz646.

8. Ingolia, N.T., Ghaemmaghami, S., Newman, J.R., and Weissman, J.S. (2009). Genome-wide analysis in vivo of translation with nucleotide resolution using ribosome profiling. Science 324, 218–223. 10.1126/science.1168978.

9. Mao, Y., Jia, L., Dong, L., Shu, X.E., and Qian, S.B. (2023). Start codon-associated ribosomal frameshifting mediates nutrient stress adaptation. Nature Structural & Molecular Biology 30, 1816–1825. 10.1038/s41594-023-01119-z.

10. Ferguson, L., Upton, H.E., Pimentel, S.C., Mok, A., Lareau, L.F., Collins, K., and Ingolia, N.T. (2023). Streamlined and sensitive mono- and di-ribosome profiling in yeast and human cells. Nature Methods 20, 1704–1715. 10.1038/s41592-023-02028-1.

11. Wang, Y., Zhang, H., and Lu, J. (2020). Recent advances in ribosome profiling for deciphering translational regulation. Methods 176, 46–54. 10.1016/j.ymeth.2019.05.011.

12. Cao, X., Sun, S., and Xing, J. (2024). A Massive Proteogenomic Screen Identifies Thousands of Novel Peptides From the Human “Dark” Proteome. Molecular & Cellular Proteomics 23, 100719. 10.1016/j.mcpro.2024.100719.

13. Ouspenskaia, T., Law, T., Clauser, K.R., Klaeger, S., Sarkizova, S., Aguet, F., Li, B., Christian, E., Knisbacher, B.A., Le, P.M., et al. (2021). Unannotated proteins expand the MHC-I-restricted immunopeptidome in cancer. Nature Biotechnology 40, 209–217. 10.1038/s41587-021-01021-3.

14. Wu, Q., Wright, M., Gogol, M.M., Bradford, W.D., Zhang, N., and Bazzini, A.A. (2020). Translation of small downstream ORFs enhances translation of canonical main open reading frames. EMBO Journal 39, e104763. 10.15252/embj.2020104763.

15. Li, Y., Zhou, H., Chen, X., Zheng, Y., Kang, Q., Hao, D., Zhang, L., Song, T., Luo, H., Hao, Y., et al. (2021). SmProt: A Reliable Repository with Comprehensive Annotation of Small Proteins Identified from Ribosome Profiling. Genomics, Proteomics & Bioinformatics 19, 602–610. 10.1016/j.gpb.2021.09.002.

16. Leblanc, S., Yala, F., Provencher, N., Lucier, J.F., Levesque, M., Lapointe, X., Jacques, J.F., Fournier, I., Salzet, M., Ouangraoua, A., et al. (2024). OpenProt 2.0 builds a path to the functional characterization of alternative proteins. Nucleic Acids Research 52, D522–d528. 10.1093/nar/gkad1050.

17. Yang, H., Li, Q., Stroup, E.K., Wang, S., and Ji, Z. (2024). Widespread stable noncanonical peptides identified by integrated analyses of ribosome profiling and ORF features. Nat Commun 15, 1932. 10.1038/s41467-024-46240-9.

18. Prensner, J.R., Abelin, J.G., Kok, L.W., Clauser, K.R., Mudge, J.M., Ruiz-Orera, J., Bassani-Sternberg, M., Moritz, R.L., Deutsch, E.W., and van Heesch, S. (2023). What Can Ribo-Seq, Immunopeptidomics, and Proteomics Tell Us About the Noncanonical Proteome? Molecular & Cellular Proteomics 22, 100631. 10.1016/j.mcpro.2023.100631.

19. Valdivia-Francia, F., and Sendoel, A. (2024). No country for old methods: New tools for studying microproteins. iScience 27, 108972. 10.1016/j.isci.2024.108972.

20. Lei, T., Chang, Y., Yao, C., and Zhang, H. (2023). A systematic evaluation of computational methods for predicting translated non-canonical ORFs from ribosome profiling data. Journal of Genetics and Genomics 51, 105–108. 10.1016/j.jgg.2023.08.010.

21. Olexiouk, V., Crappé, J., Verbruggen, S., Verhegen, K., Martens, L., and Menschaert, G. (2016). sORFs.org: a repository of small ORFs identified by ribosome profiling. Nucleic Acids Research 44, D324–D329. 10.1093/nar/gkv1175.

22. Chothani, S.P., Adami, E., Widjaja, A.A., Langley, S.R., Viswanathan, S., Pua, C.J., Zhihao, N.T., Harmston, N., D’Agostino, G., Whiffin, N., et al. (2022). A high-resolution map of human RNA translation. Molecular Cell 82, 2885–2899.e2888. 10.1016/j.molcel.2022.06.023.

23. Prensner, J.R., Enache, O.M., Luria, V., Krug, K., Clauser, K.R., Dempster, J.M., Karger, A., Wang, L., Stumbraite, K., Wang, V.M., et al. (2021). Noncanonical open reading frames encode functional proteins essential for cancer cell survival. Nature Biotechnology 39, 697–704. 10.1038/s41587-020-00806-2.

24. Camarena, M.E., Theunissen, P., Ruiz, M., Ruiz-Orera, J., Calvo-Serra, B., Castelo, R., Castro, C., Sarobe, P., Fortes, P., Perera-Bel, J., and Albà, M.M. (2024). Microproteins encoded by noncanonical ORFs are a major source of tumor-specific antigens in a liver cancer patient meta-cohort. Sci Adv 10, eadn3628. 10.1126/sciadv.adn3628.

25. Sendoel, A., Dunn, J.G., Rodriguez, E.H., Naik, S., Gomez, N.C., Hurwitz, B., Levorse, J., Dill, B.D., Schramek, D., Molina, H., et al. (2017). Translation from unconventional 5’ start sites drives tumour initiation. Nature 541, 494–499. 10.1038/nature21036.

26. Robichaud, N., Sonenberg, N., Ruggero, D., and Schneider, R.J. (2019). Translational Control in Cancer. Cold Spring Harb Perspect Biol 11. 10.1101/cshperspect.a032896.

27. Malekos, E., and Carpenter, S. (2022). Short open reading frame genes in innate immunity: from discovery to characterization. Trends Immunol 43, 741–756. 10.1016/j.it.2022.07.005.

28. Chen, J., Brunner, A.D., Cogan, J.Z., Nunez, J.K., Fields, A.P., Adamson, B., Itzhak, D.N., Li, J.Y., Mann, M., Leonetti, M.D., and Weissman, J.S. (2020). Pervasive functional translation of noncanonical human open reading frames. Science 367, 1140–1146. 10.1126/science.aay0262.

29. Zheng, C., Wei, Y., Zhang, P., Lin, K., He, D., Teng, H., Manyam, G., Zhang, Z., Liu, W., Lee, H.R.L., et al. (2023). CRISPR-Cas9-based functional interrogation of unconventional translatome reveals human cancer dependency on cryptic non-canonical open reading frames. Nature Structural & Molecular Biology 30, 1878–1892. 10.1038/s41594-023-01117-1.

30. Zheng, C., Wei, Y., Zhang, P., Xu, L., Zhang, Z., Lin, K., Hou, J., Lv, X., Ding, Y., Chiu, Y., et al. (2023). CRISPR/Cas9 screen uncovers functional translation of cryptic lncRNA-encoded open reading frames in human cancer. J Clin Invest 133. 10.1172/jci159940.

31. Schlesinger, D., Dirks, C., Luzon, C.N., Lafranchi, L., Eirich, J., and Elsässer, S.J. (2023). A large-scale sORF screen identifies putative microproteins and provides insights into their interaction partners, localisation and function. bioRxiv, 2023.2006.2013.544808. 10.1101/2023.06.13.544808.

32. Hofman, D.A., Ruiz-Orera, J., Yannuzzi, I., Murugesan, R., Brown, A., Clauser, K.R., Condurat, A.L., van Dinter, J.T., Engels, S.A.G., Goodale, A., et al. (2024). Translation of non-canonical open reading frames as a cancer cell survival mechanism in childhood medulloblastoma. Molecular Cell 84, 261–276.e218. 10.1016/j.molcel.2023.12.003.

33. Vakirlis, N., Vance, Z., Duggan, K.M., and McLysaght, A. (2022). De novo birth of functional microproteins in the human lineage. Cell Rep 41, 111808. 10.1016/j.celrep.2022.111808.

34. Sandmann, C.L., Schulz, J.F., Ruiz-Orera, J., Kirchner, M., Ziehm, M., Adami, E., Marczenke, M., Christ, A., Liebe, N., Greiner, J., et al. (2023). Evolutionary origins and interactomes of human, young microproteins and small peptides translated from short open reading frames. Mol Cell 83, 994–1011.e1018. 10.1016/j.molcel.2023.01.023.

35. Wacholder, A., Parikh, S.B., Coelho, N.C., Acar, O., Houghton, C., Chou, L., and Carvunis, A.R. (2023). A vast evolutionarily transient translatome contributes to phenotype and fitness. Cell Syst. 10.1016/j.cels.2023.04.002.

36. Erhard, F., Halenius, A., Zimmermann, C., L’Hernault, A., Kowalewski, D.J., Weekes, M.P., Stevanovic, S., Zimmer, R., and Dolken, L. (2018). Improved Ribo-seq enables identification of cryptic translation events. Nat Methods 15, 363–366. 10.1038/nmeth.4631.

37. Xiao, Z., Huang, R., Xing, X., Chen, Y., Deng, H., and Yang, X. (2018). De novo annotation and characterization of the translatome with ribosome profiling data. Nucleic Acids Res 46, e61. 10.1093/nar/gky179.

38. Zhang, P., He, D., Xu, Y., Hou, J., Pan, B.F., Wang, Y., Liu, T., Davis, C.M., Ehli, E.A., Tan, L., et al. (2017). Genome-wide identification and differential analysis of translational initiation. Nat Commun 8, 1749. 10.1038/s41467-017-01981-8.

39. Ingolia, N.T., Brar, G.A., Stern-Ginossar, N., Harris, M.S., Talhouarne, G.J., Jackson, S.E., Wills, M.R., and Weissman, J.S. (2014). Ribosome profiling reveals pervasive translation outside of annotated protein-coding genes. Cell Rep 8, 1365–1379. 10.1016/j.celrep.2014.07.045.

40. Chew, G.L., Pauli, A., Rinn, J.L., Regev, A., Schier, A.F., and Valen, E. (2013). Ribosome profiling reveals resemblance between long non-coding RNAs and 5’ leaders of coding RNAs. Development 140, 2828–2834. 10.1242/dev.098343.

41. Guttman, M., Russell, P., Ingolia, N.T., Weissman, J.S., and Lander, E.S. (2013). Ribosome profiling provides evidence that large noncoding RNAs do not encode proteins. Cell 154, 240–251. 10.1016/j.cell.2013.06.009.

42. Morales, J., Pujar, S., Loveland, J.E., Astashyn, A., Bennett, R., Berry, A., Cox, E., Davidson, C., Ermolaeva, O., Farrell, C.M., et al. (2022). A joint NCBI and EMBL-EBI transcript set for clinical genomics and research. Nature 604, 310–315. 10.1038/s41586-022-04558-8.

43. Pozo, F., Rodriguez, J.M., Martínez Gómez, L., Vázquez, J., and Tress, M.L. (2022). APPRIS principal isoforms and MANE Select transcripts define reference splice variants. Bioinformatics 38, ii89–ii94. 10.1093/bioinformatics/btac473.

44. Chothani, S., Ruiz-Orera, J., Tierney, J.A.S., Clauwaert, J., Deutsch, E.W., Alba, M.M., Aspden, J.L., Baranov, P.V., Bazzini, A.A., Bruford, E.A., et al. (2025). An expanded reference catalog of translated open reading frames for biomedical research. bioRxiv. 10.1101/2025.07.03.662928.

45. Deutsch, E.W., Kok, L.W., Mudge, J.M., Ruiz-Orera, J., Fierro-Monti, I., Sun, Z., Abelin, J.G., Alba, M.M., Aspden, J.L., Bazzini, A.A., et al. (2024). High-quality peptide evidence for annotating non-canonical open reading frames as human proteins. bioRxiv. 10.1101/2024.09.09.612016.

46. Shabalina, S.A., Spiridonov, N.A., and Kashina, A. (2013). Sounds of silence: synonymous nucleotides as a key to biological regulation and complexity. Nucleic Acids Res 41, 2073–2094. 10.1093/nar/gks1205.

47. Hernández, G., Osnaya, V.G., and Pérez-Martínez, X. (2019). Conservation and Variability of the AUG Initiation Codon Context in Eukaryotes. Trends Biochem Sci 44, 1009–1021. 10.1016/j.tibs.2019.07.001.

48. Zhang, H., Wang, Y., Wu, X., Tang, X., Wu, C., and Lu, J. (2021). Determinants of genome-wide distribution and evolution of uORFs in eukaryotes. Nature Communications 12, 1076. 10.1038/s41467-021-21394-y.

49. Wells, J.N., and Feschotte, C. (2020). A Field Guide to Eukaryotic Transposable Elements. Annu Rev Genet 54, 539–561. 10.1146/annurev-genet-040620-022145.

50. Chen, S., Francioli, L.C., Goodrich, J.K., Collins, R.L., Kanai, M., Wang, Q., Alföldi, J., Watts, N.A., Vittal, C., Gauthier, L.D., et al. (2024). A genomic mutational constraint map using variation in 76,156 human genomes. Nature 625, 92–100. 10.1038/s41586-023-06045-0.

51. Kuderna, L.F.K., Ulirsch, J.C., Rashid, S., Ameen, M., Sundaram, L., Hickey, G., Cox, A.J., Gao, H., Kumar, A., Aguet, F., et al. (2024). Identification of constrained sequence elements across 239 primate genomes. Nature 625, 735–742. 10.1038/s41586-023-06798-8.

52. Pollard, K.S., Hubisz, M.J., Rosenbloom, K.R., and Siepel, A. (2010). Detection of nonneutral substitution rates on mammalian phylogenies. Genome Res 20, 110–121. 10.1101/gr.097857.109.

53. Lin, M.F., Jungreis, I., and Kellis, M. (2011). PhyloCSF: a comparative genomics method to distinguish protein coding and non-coding regions. Bioinformatics 27, i275–282. 10.1093/bioinformatics/btr209.

54. Stark, A., Lin, M.F., Kheradpour, P., Pedersen, J.S., Parts, L., Carlson, J.W., Crosby, M.A., Rasmussen, M.D., Roy, S., Deoras, A.N., et al. (2007). Discovery of functional elements in 12 Drosophila genomes using evolutionary signatures. Nature 450, 219–232. 10.1038/nature06340.

55. Wang, D., Eraslan, B., Wieland, T., Hallstrom, B., Hopf, T., Zolg, D.P., Zecha, J., Asplund, A., Li, L.H., Meng, C., et al. (2019). A deep proteome and transcriptome abundance atlas of 29 healthy human tissues. Mol Syst Biol 15, e8503. 10.15252/msb.20188503.

56. Giansanti, P., Samaras, P., Bian, Y., Meng, C., Coluccio, A., Frejno, M., Jakubowsky, H., Dobiasch, S., Hazarika, R.R., Rechenberger, J., et al. (2022). Mass spectrometry-based draft of the mouse proteome. Nat Methods 19, 803–811. 10.1038/s41592-022-01526-y.

57. Phillips, P.C. (2008). Epistasis--the essential role of gene interactions in the structure and evolution of genetic systems. Nat Rev Genet 9, 855–867. 10.1038/nrg2452.

58. Langfelder, P., and Horvath, S. (2008). WGCNA: an R package for weighted correlation network analysis. BMC Bioinformatics 9, 559. 10.1186/1471-2105-9-559.

59. Zhang, H., Dou, S., He, F., Luo, J., Wei, L., and Lu, J. (2018). Genome-wide maps of ribosomal occupancy provide insights into adaptive evolution and regulatory roles of uORFs during Drosophila development. PLoS Biology 16, e2003903. 10.1371/journal.pbio.2003903.

60. Chew, G.L., Pauli, A., and Schier, A.F. (2016). Conservation of uORF repressiveness and sequence features in mouse, human and zebrafish. Nat Commun 7, 11663. 10.1038/ncomms11663.

61. Zhang, J., and Xu, C. (2022). Gene product diversity: adaptive or not? Trends Genet 38, 1112–1122. 10.1016/j.tig.2022.05.002.

62. Mao, Y., and Qian, S.B. (2024). Making sense of mRNA translational “noise”. Semin Cell Dev Biol 154, 114–122. 10.1016/j.semcdb.2023.03.004.

63. Broeils, L.A., Ruiz-Orera, J., Snel, B., Hubner, N., and van Heesch, S. (2023). Evolution and implications of de novo genes in humans. Nat Ecol Evol. 10.1038/s41559-023-02014-y.

64. Liu, X., Xiao, C., Xu, X., Zhang, J., Mo, F., Chen, J.Y., Delihas, N., Zhang, L., An, N.A., and Li, C.Y. (2024). Origin of functional de novo genes in humans from “hopeful monsters”. Wiley Interdiscip Rev RNA 15, e1845. 10.1002/wrna.1845.

65. Mackowiak, S.D., Zauber, H., Bielow, C., Thiel, D., Kutz, K., Calviello, L., Mastrobuoni, G., Rajewsky, N., Kempa, S., Selbach, M., and Obermayer, B. (2015). Extensive identification and analysis of conserved small ORFs in animals. Genome Biology 16, 179. 10.1186/s13059-015-0742-x.

66. Sun, Y., Duan, Y., Gao, P., Liu, C., Jin, K., Dou, S., Tang, W., Zhang, H., and Lu, J. (2025). Upstream open reading frames buffer translational variability during Drosophila evolution and development. Elife 14. 10.7554/eLife.104074.

67. Treichel, A.J., and Bazzini, A.A. (2022). Casting CRISPR-Cas13d to fish for microprotein functions in animal development. iScience 25, 105547. 10.1016/j.isci.2022.105547.

68. Martin, M. (2011). Cutadapt removes adapter sequences from high-throughput sequencing reads. EMBnet. journal 17, 10–12.

69. Cunningham, F., Allen, J.E., Allen, J., Alvarez-Jarreta, J., Amode, M.R., Armean, I.M., Austine-Orimoloye, O., Azov, A.G., Barnes, I., Bennett, R., et al. (2022). Ensembl 2022. Nucleic Acids Res 50, D988–d995. 10.1093/nar/gkab1049.

70. Dobin, A., Davis, C.A., Schlesinger, F., Drenkow, J., Zaleski, C., Jha, S., Batut, P., Chaisson, M., and Gingeras, T.R. (2013). STAR: ultrafast universal RNA-seq aligner. Bioinformatics 29, 15–21. 10.1093/bioinformatics/bts635.

71. Chang, Y., Lei, T., and Zhang, H. (2023). PSite: inference of read-specific P-site offsets for ribosomal footprints. bioRxiv, 2023.2006.2027.546788. 10.1101/2023.06.27.546788.

72. Tierney, J.A.S., Świrski, M.I., Tjeldnes, H., Kiran, A.M., Carancini, G., Kiniry, S.J., Michel, A.M., Kufel, J., Valen, E., and Baranov, P.V. (2025). RiboSeq.Org: an integrated suite of resources for ribosome profiling data analysis and visualization. Nucleic Acids Res 53, D268–d274. 10.1093/nar/gkae1020.

73. Nassar, L.R., Barber, G.P., Benet-Pagès, A., Casper, J., Clawson, H., Diekhans, M., Fischer, C., Gonzalez, J.N., Hinrichs, A.S., Lee, B.T., et al. (2023). The UCSC Genome Browser database: 2023 update. Nucleic Acids Res 51, D1188–d1195. 10.1093/nar/gkac1072.

74. Rodriguez, J.M., Pozo, F., Cerdán-Vélez, D., Di Domenico, T., Vázquez, J., and Tress, M.L. (2022). APPRIS: selecting functionally important isoforms. Nucleic Acids Res 50, D54–d59. 10.1093/nar/gkab1058.

75. Lorenz, R., Bernhart, S.H., Höner Zu Siederdissen, C., Tafer, H., Flamm, C., Stadler, P.F., and Hofacker, I.L. (2011). ViennaRNA Package 2.0. Algorithms Mol Biol 6, 26. 10.1186/1748-7188-6-26.

76. Liu, M., Zi, B., Zhang, H., and Zhang, H. (2024). cubar: a versatile package for codon usage bias analysis in R. R package version 1.1. 10.5281/zenodo.10155990.

77. Chan, P.P., and Lowe, T.M. (2016). GtRNAdb 2.0: an expanded database of transfer RNA genes identified in complete and draft genomes. Nucleic Acids Res 44, D184–189. 10.1093/nar/gkv1309.

78. Jones, P., Binns, D., Chang, H.Y., Fraser, M., Li, W., McAnulla, C., McWilliam, H., Maslen, J., Mitchell, A., Nuka, G., et al. (2014). InterProScan 5: genome-scale protein function classification. Bioinformatics 30, 1236–1240. 10.1093/bioinformatics/btu031.

79. Quinlan, A.R., and Hall, I.M. (2010). BEDTools: a flexible suite of utilities for comparing genomic features. Bioinformatics 26, 841–842. 10.1093/bioinformatics/btq033.

80. Erdős, G., Pajkos, M., and Dosztányi, Z. (2021). IUPred3: prediction of protein disorder enhanced with unambiguous experimental annotation and visualization of evolutionary conservation. Nucleic Acids Res 49, W297–w303. 10.1093/nar/gkab408.

81. Dalla, V., Giraitis, L., and Phillips, P.C.B. (2022). obust Tests for White Noise and Cross-Correlation. Econometric Theory 38, 913–941. 10.1017/S0266466620000341.

82. Katoh, K., and Standley, D.M. (2013). MAFFT multiple sequence alignment software version 7: improvements in performance and usability. Mol Biol Evol 30, 772–780. 10.1093/molbev/mst010.

83. Lemoine, F., and Gascuel, O. (2021). Gotree/Goalign: toolkit and Go API to facilitate the development of phylogenetic workflows. NAR Genom Bioinform 3, lqab075. 10.1093/nargab/lqab075.

84. Ashkenazy, H., Penn, O., Doron-Faigenboim, A., Cohen, O., Cannarozzi, G., Zomer, O., and Pupko, T. (2012). FastML: a web server for probabilistic reconstruction of ancestral sequences. Nucleic Acids Res 40, W580–584. 10.1093/nar/gks498.

85. Paradis, E., and Schliep, K. (2019). ape 5.0: an environment for modern phylogenetics and evolutionary analyses in R. Bioinformatics 35, 526–528. 10.1093/bioinformatics/bty633.

86. Pockrandt, C., Steinegger, M., and Salzberg, S.L. (2022). PhyloCSF++: a fast and user-friendly implementation of PhyloCSF with annotation tools. Bioinformatics 38, 1440–1442. 10.1093/bioinformatics/btab756.

87. Love, M.I., Huber, W., and Anders, S. (2014). Moderated estimation of fold change and dispersion for RNA-seq data with DESeq2. Genome Biol 15, 550. 10.1186/s13059-014-0550-8.

88. Kryuchkova-Mostacci, N., and Robinson-Rechavi, M. (2017). A benchmark of gene expression tissue-specificity metrics. Brief Bioinform 18, 205–214. 10.1093/bib/bbw008.

89. Patro, R., Duggal, G., Love, M.I., Irizarry, R.A., and Kingsford, C. (2017). Salmon provides fast and bias-aware quantification of transcript expression. Nat Methods 14, 417–419. 10.1038/nmeth.4197.

90. Rezaie, N., Reese, F., and Mortazavi, A. (2023). PyWGCNA: a Python package for weighted gene co-expression network analysis. Bioinformatics 39. 10.1093/bioinformatics/btad415.

91. Traag, V.A., Waltman, L., and van Eck, N.J. (2019). From Louvain to Leiden: guaranteeing well-connected communities. Sci Rep 9, 5233. 10.1038/s41598-019-41695-z.

92. Csárdi, G., Nepusz, T., Traag, V., Horvát, S., Zanini, F., Noom, D., and Müller, K. (2024). igraph: Network Analysis and Visualization in R. R package version 2.1.3. 10.5281/zenodo.7682609.

93. Wu, T., Hu, E., Xu, S., Chen, M., Guo, P., Dai, Z., Feng, T., Zhou, L., Tang, W., Zhan, L., et al. (2021). clusterProfiler 4.0: A universal enrichment tool for interpreting omics data. Innovation (Camb) 2, 100141. 10.1016/j.xinn.2021.100141.

