## Supplementary Information for "Evolutionary remodeling of non-canonical ORF translation in mammals"

### Supplementary Figures

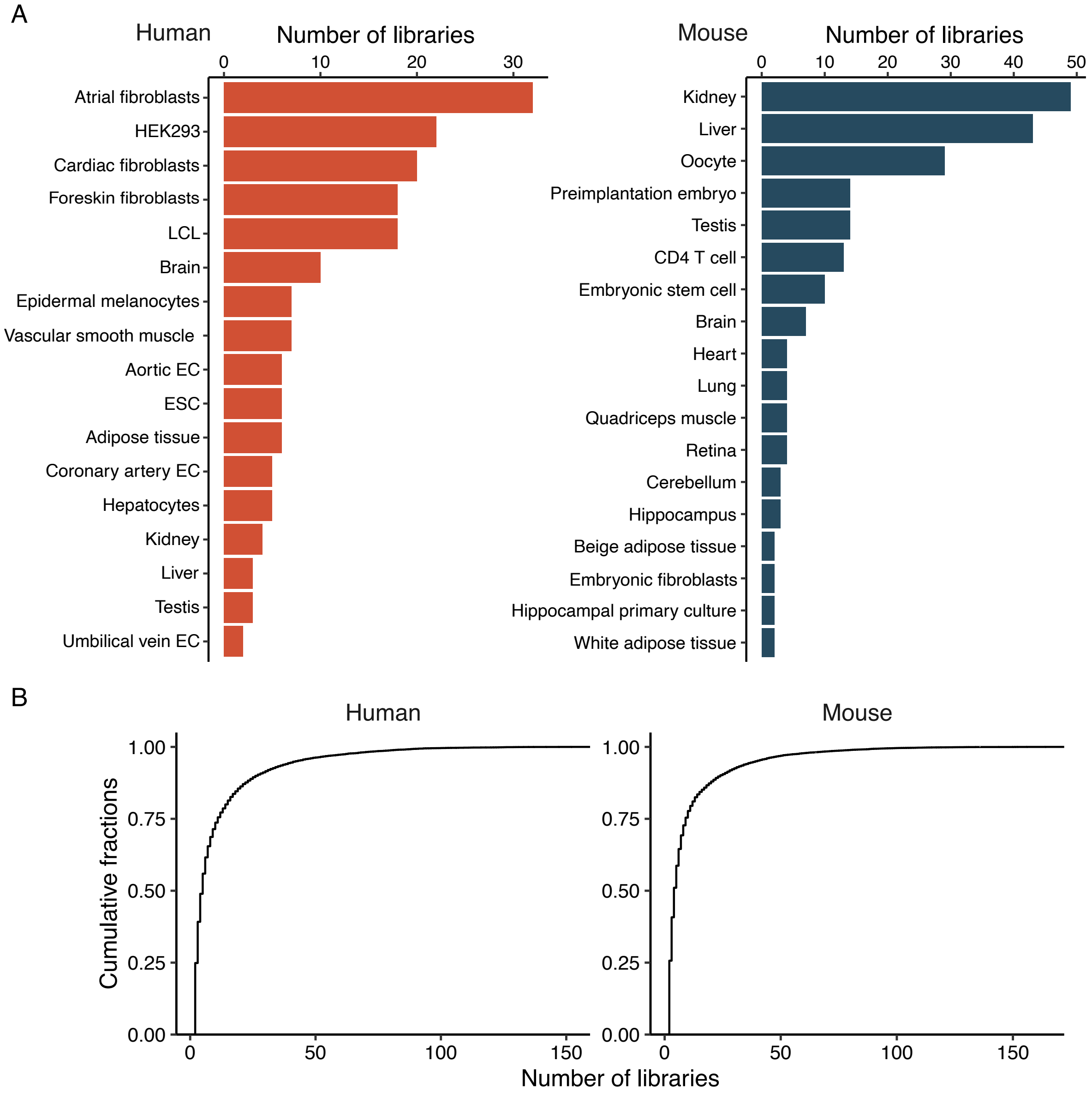


**Supplementary figure 1.** Overview of ncORF annotation across different libraries. (A) Number of high-quality Ribo-Seq libraries passing quality control for each sample in humans and mice. (B) Cumulative faction of ncORFs detected across different numbers of Ribo-Seq libraries in humans and mice. EC, endothelial cell; ESC, embryonic stem cell; LCL, lymphoblastoid cell line; Ribo-Seq, ribosome profiling.

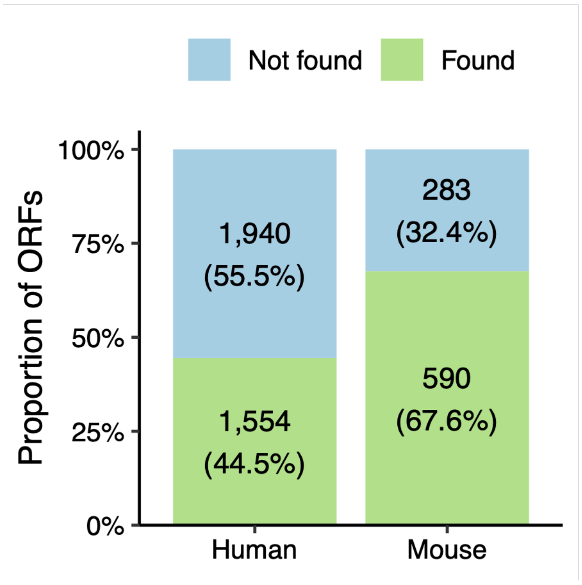


**Supplementary figure 2.** Rediscovery rate of non-canonical ORFs supported by MS evidence, as compiled in our previous study[1]. MS, mass spectrometry; ORF, open reading frame.

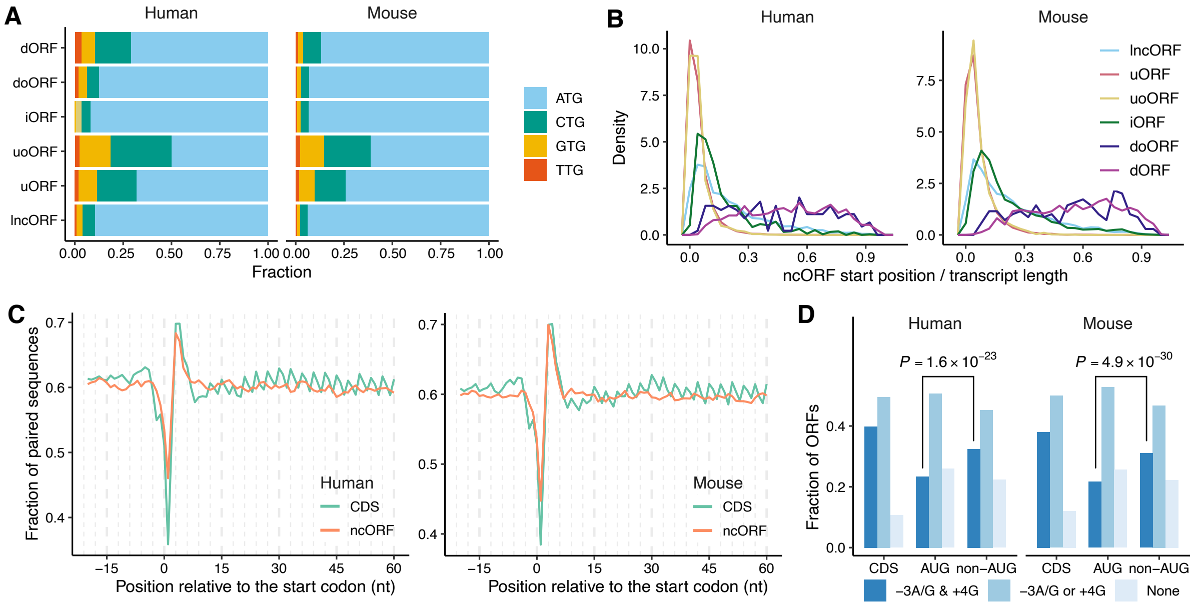


**Supplementary figure 3.** Translation-related signatures of ncORF sequences. (**A**) Start codon usage among ncORFs. (**B**) Distribution of relative start positions of ncORFs within transcripts. The *x*-axis represents the position of ncORF start codon normalized by transcript length. (**C**) Nucleotide pairing probabilities around the start codons of ncORFs and CDSs. (**D**) Proportion of ncORFs with different Kozak sequence contexts. Statistical significance was assessed using Fisher's exact test. CDS, coding sequence; ncORF, non-canonical open reading frame; ORF, open reading frame.

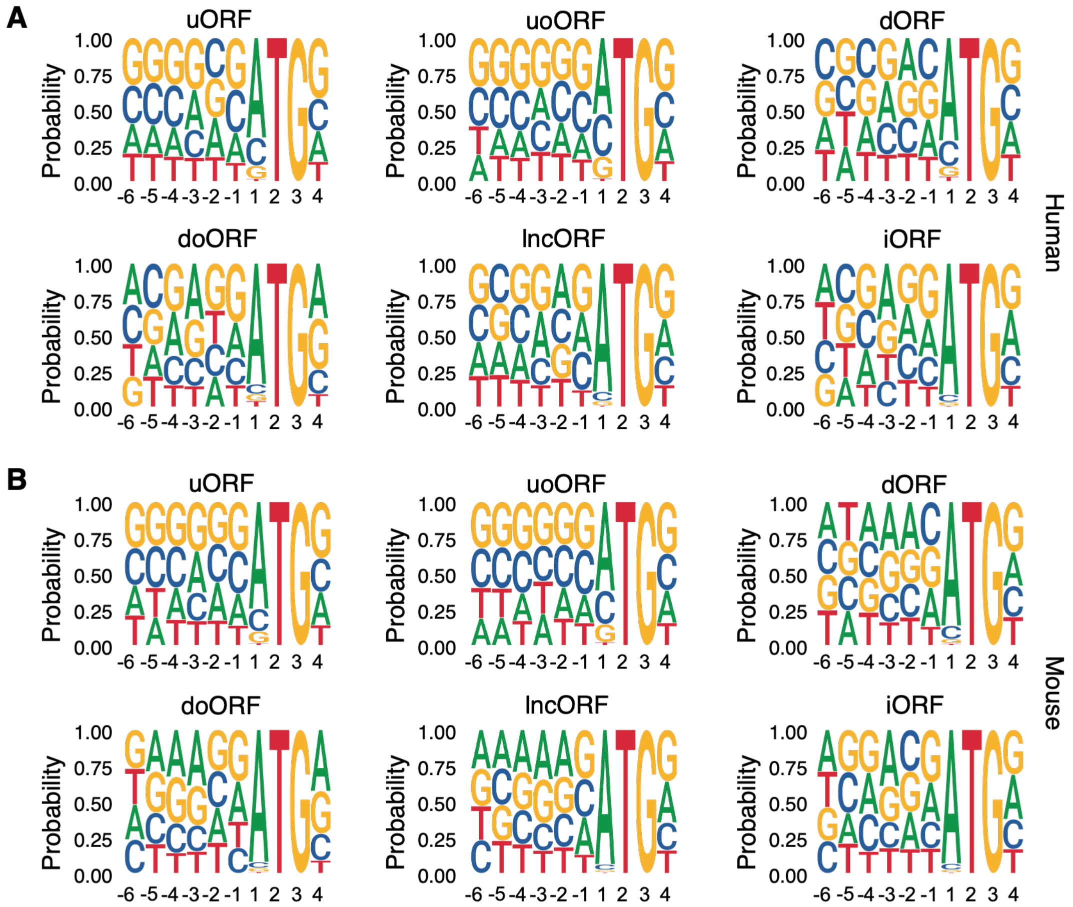


**Supplementary figure 4.** Seqlogo plots of Kozak sequence context surrounding ncORF start codons in humans (A) and mice (B). The first nucleotide of the start codon is designated as position +1. The height of each nucleotide at each position reflects its frequency. ncORF, non-canonical open reading frame.

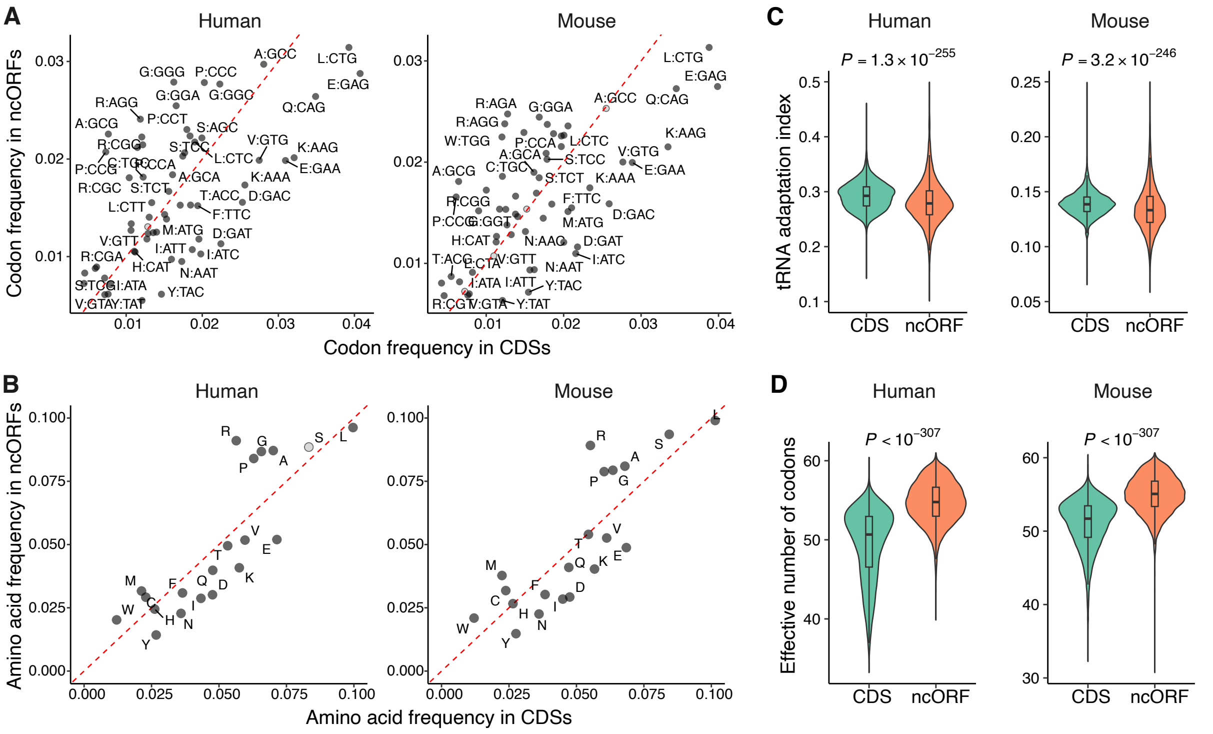


**Supplementary figure 5.** Differences in codon and amino acid usage between ncORFs and CDSs. **(A-B)** Differences in relative usage of codons (A) and amino acids (B). Significance was determined using Fisher's exact tests and corrected for multiple-testing. Codons or amino acids with adjusted *P* value < 0.05 are shown in dark grey. **(C-D)** Distribution of tRNA adaptation index values (C) and effective number of codons (D) for CDSs and ncORFs. Statistical significance was determined by Wilcoxon rank-sum tests. CDS, coding sequence; ncORF, non-canonical open reading frame.

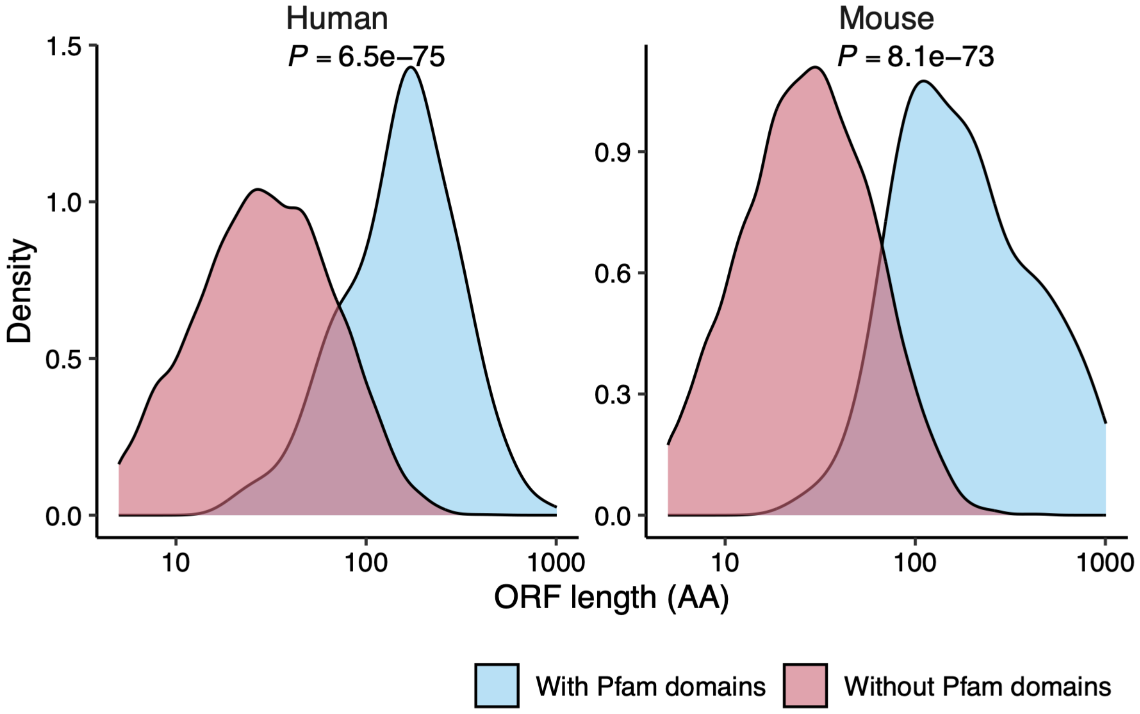


**Supplementary figure 6.** Length distribution of ncORFs with (blue) or without (red) known Pfam domains. Statistical differences were assessed using Wilcoxon rank-sum tests. AA, amino acid; ncORF, non-canonical open reading frame; ORF, open reading frame.

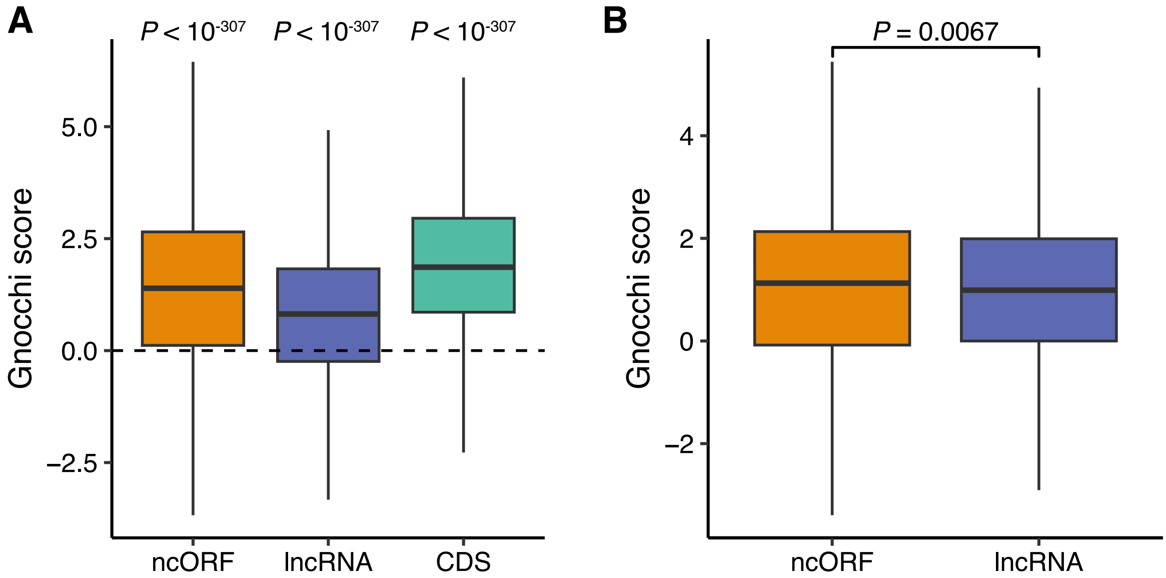


**Supplementary figure 7.** Gnocchi scores for CDSs, ncORFs, and lncRNAs. (**A**) Distribution of Gnocchi scores for all CDSs, ncORFs, and lncRNAs. ncORFs overlapping annotated CDSs were excluded. For lncRNAs, regions overlapping any CDS were removed prior to analysis. Deviation of scores from zero was assessed using two-sided Wilcoxon signed-rank tests. (**B**) Comparison of Gnocchi scores between lncRNA-derived ncORFs (lncORFs) and their corresponding full-length lncRNAs. Paired differences were evaluated using a two-sided Wilcoxon signed-rank test. CDS, coding sequence; Gnocchi, Genomic Non-Coding Constraint of Haplo-Insufficient variation; lncRNA, long non-coding RNA; ncORF, non-canonical open reading frame.

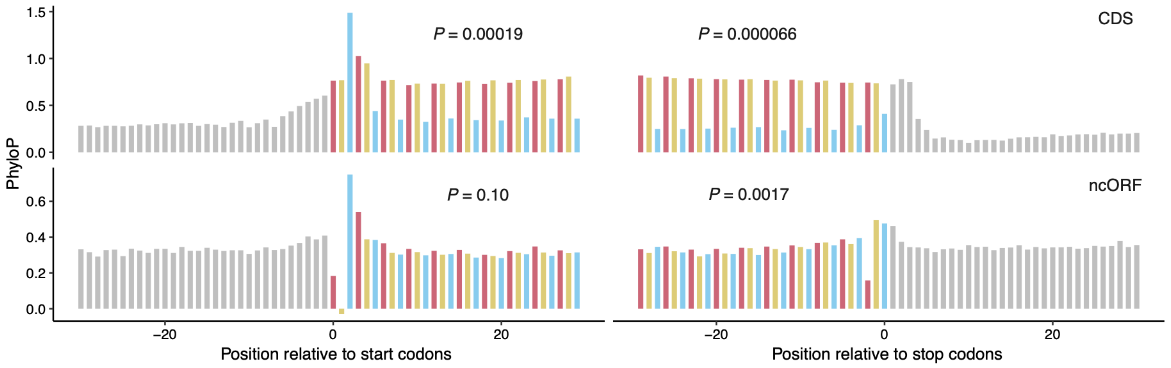


**Supplementary figure 8.** The mean PhyloP score of 30 base pairs upstream and downstream of the ORF (CDS or ncORF) start or stop codons in primates. Codons were delineated with bars of alternating colors for different frames, and nucleotides in untranslated regions were shown in grey. Statistical significance of the three-nucleotide periodicity was determined by autocorrelation with a lag of three. CDS, coding sequence; ncORF, non-canonical open reading frame.

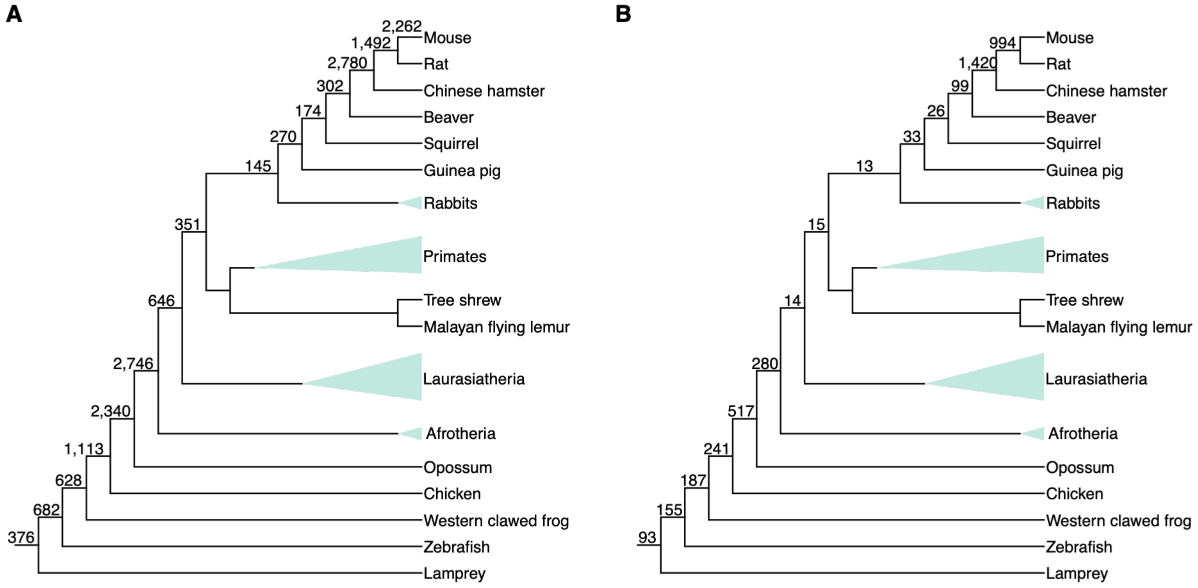


**Supplementary figure 9.** Cladograms of vertebrates showing the number of ncORFs originating at each ancestral branch leading to mice. (**A**) All ncORFs. (**B**) Lineage-specific ncORFs with local branch length score > 0.9. Triangles indicate species merged into larger clades for visual simplicity.

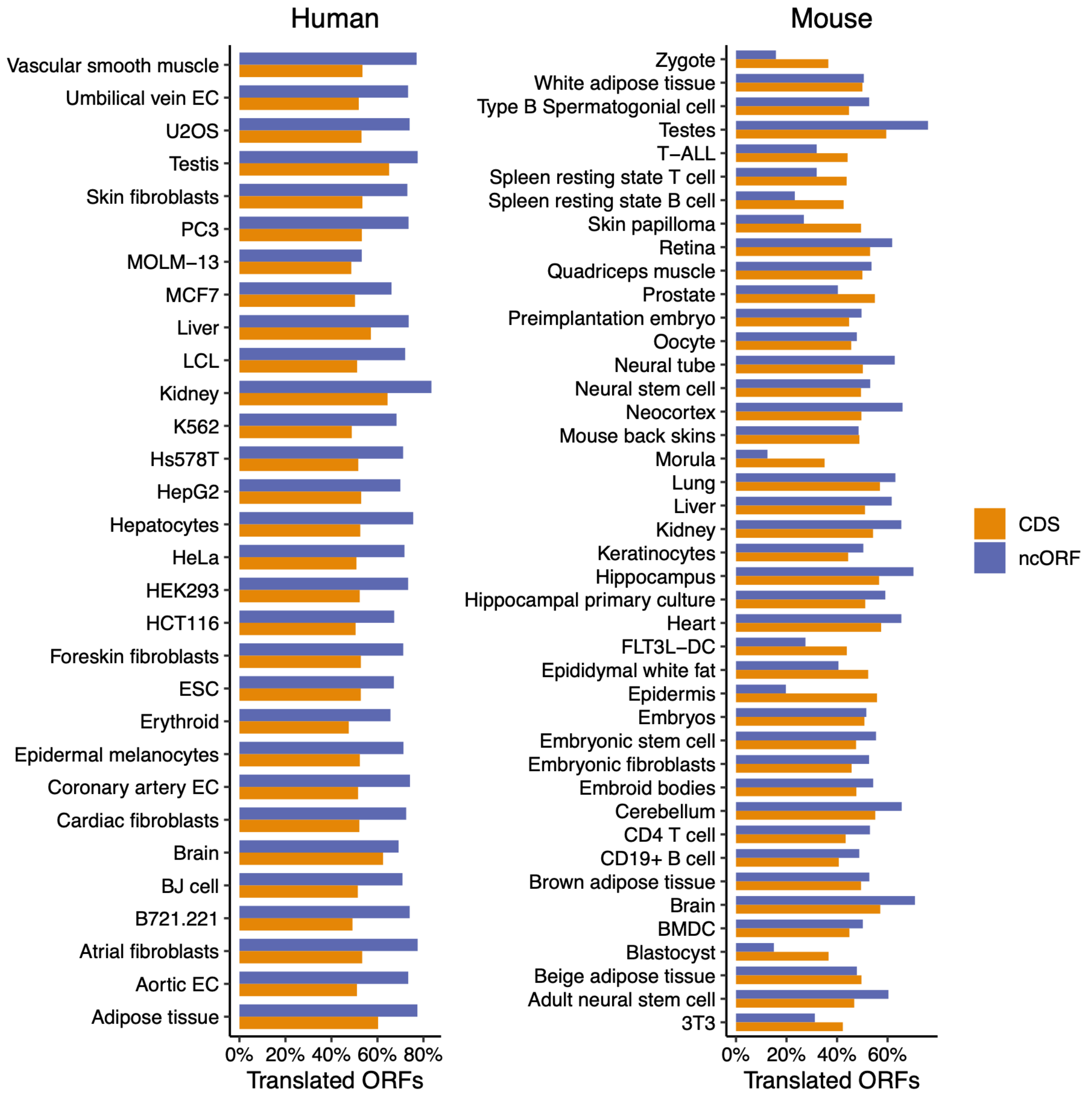


**Supplementary figure 10**. Proportion of translated ncORFs and CDSs detected in each human and mouse sample. Translated ORFs were detected with RPKM ≥ 1. BMDC, bone marrow-derived dendritic cell; CDS, coding sequence; DC, dendritic cell; EC, endothelial cell; ESC, embryonic stem cell; LCL, lymphoblastoid cell line; ncORF, non-canonical open reading frame; RPKM, reads per kilobase per million mapped reads; T-ALL, T-cell acute lymphoblastic leukemia.

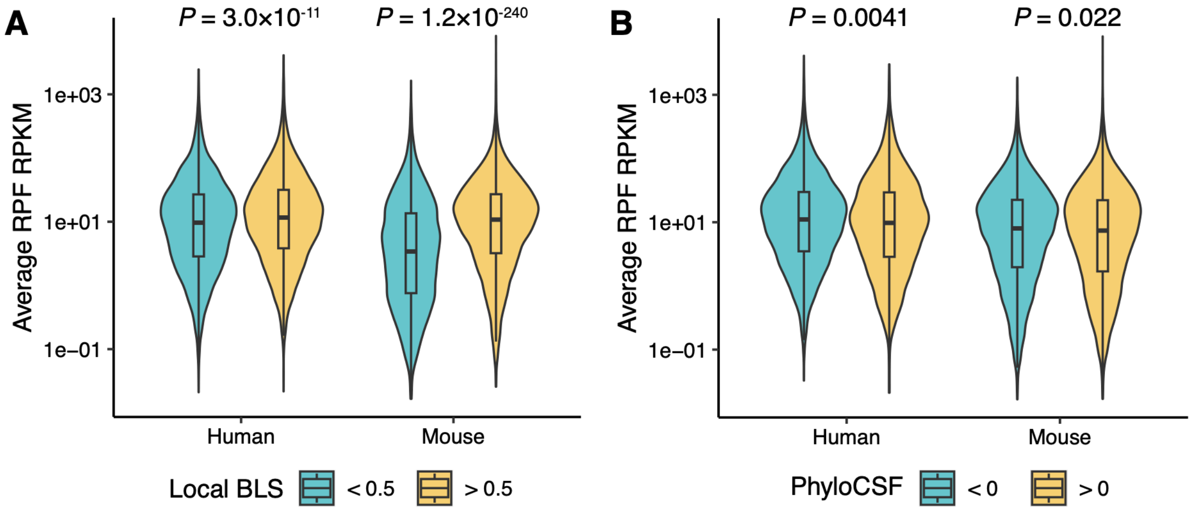


**Supplementary figure 11.** Distribution of average translation levels of ncORFs stratified by local BLS (A) or PhyloCSF (B) in humans and mice. Statistical differences were assessed using Wilcoxon rank-sum tests. BLS, branch length score; ncORF, non-canonical open reading frame; RPF, ribosome-protected fragment; RPKM, reads per kilobase per million mapped reads.

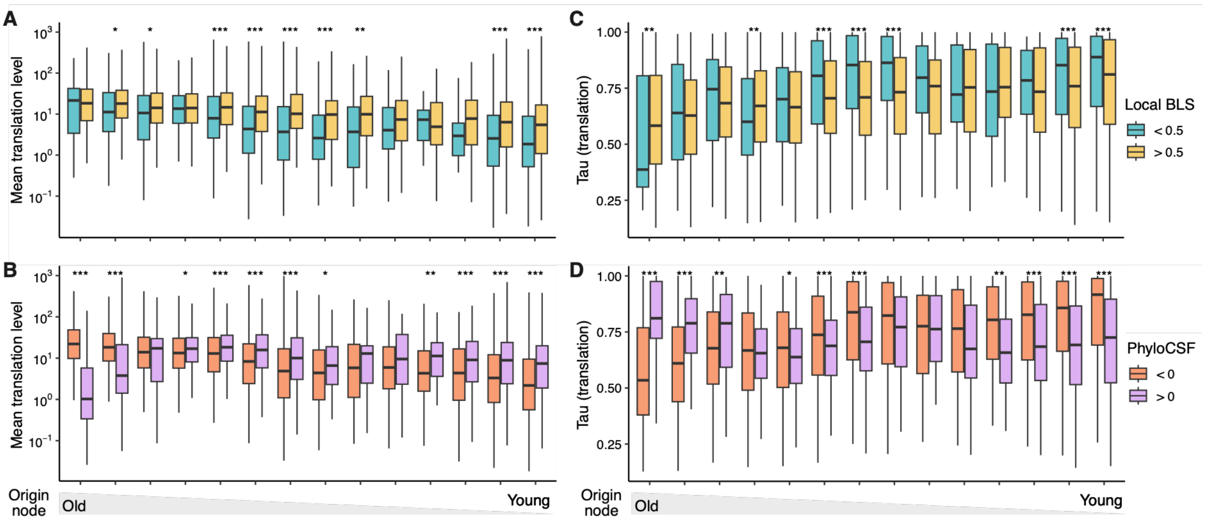


**Supplementary figure 12.** Expression patterns of ncORFs in mice. (**A**) Distribution of mean translation levels of mouse ncORFs grouped by origin nodes and further stratified by local BLS. Statistical significance was assessed with Wilcoxon rank-sum tests. ***, $P < 0.001$; **, $P < 0.01$; *, $P < 0.05$. (**B**) Similar to (A) but ncORFs are further stratified by PhyloCSF scores. (**C-D**) Similar to (A) and (B), but showing the distribution of tissue-specificity at the translation level.

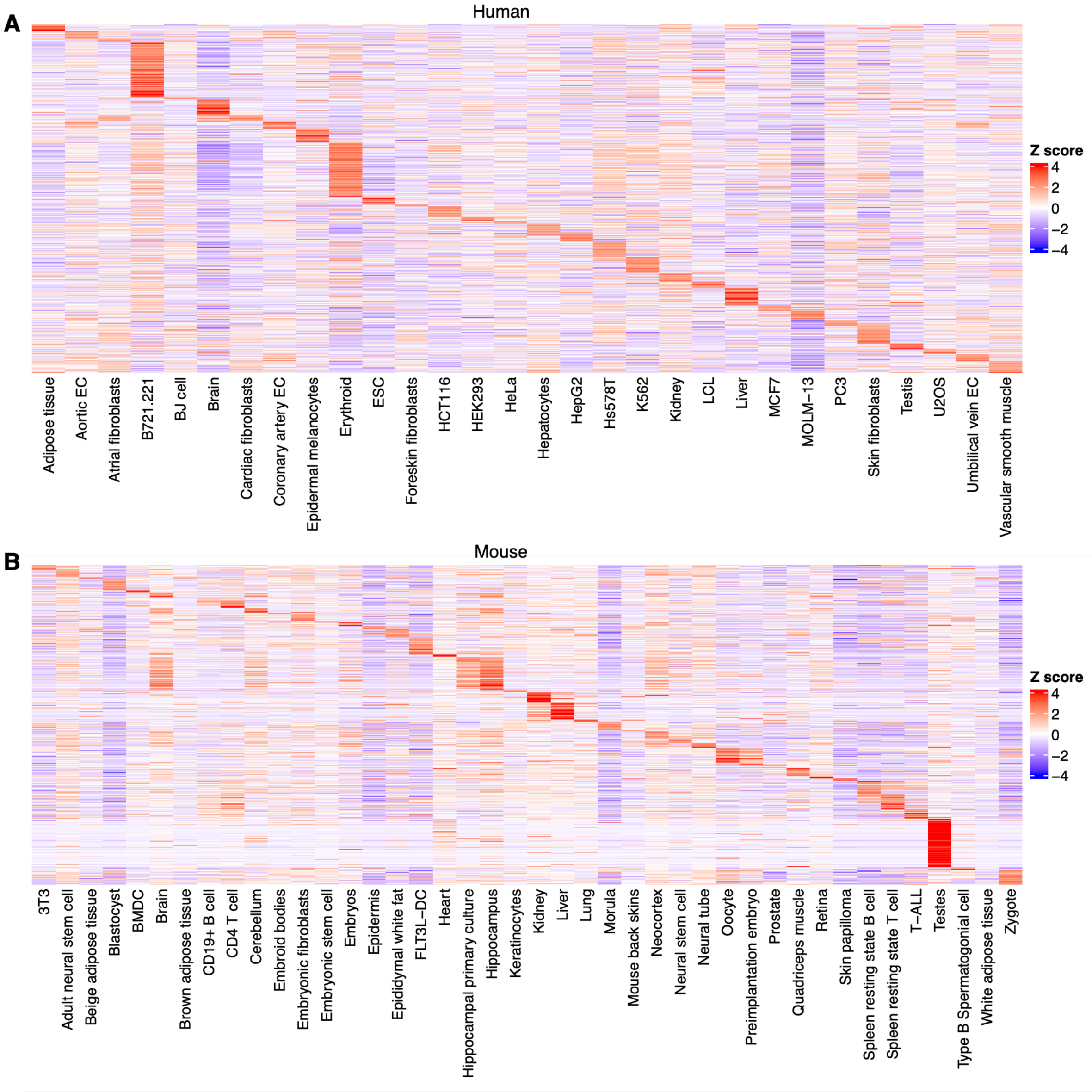


**Supplementary figure 13.** Heatmap showing the translation levels of ncORFs in human (A) and mouse (B) samples. Only ncORFs with translation level (RPKM) ≥ 1 in at least one sample are displayed. Translation levels were normalized across samples by Z-score transformation before visualization. BMDC, bone marrow-derived dendritic cell; DC, dendritic cell; EC, endothelial cell; ESC, embryonic stem cell; LCL, lymphoblastoid cell line; ncORF, non-canonical open reading frame; Ribo-Seq, ribosome profiling; RPKM, reads per kilobase per million mapped reads; T-ALL, T-cell acute lymphoblastic leukemia.

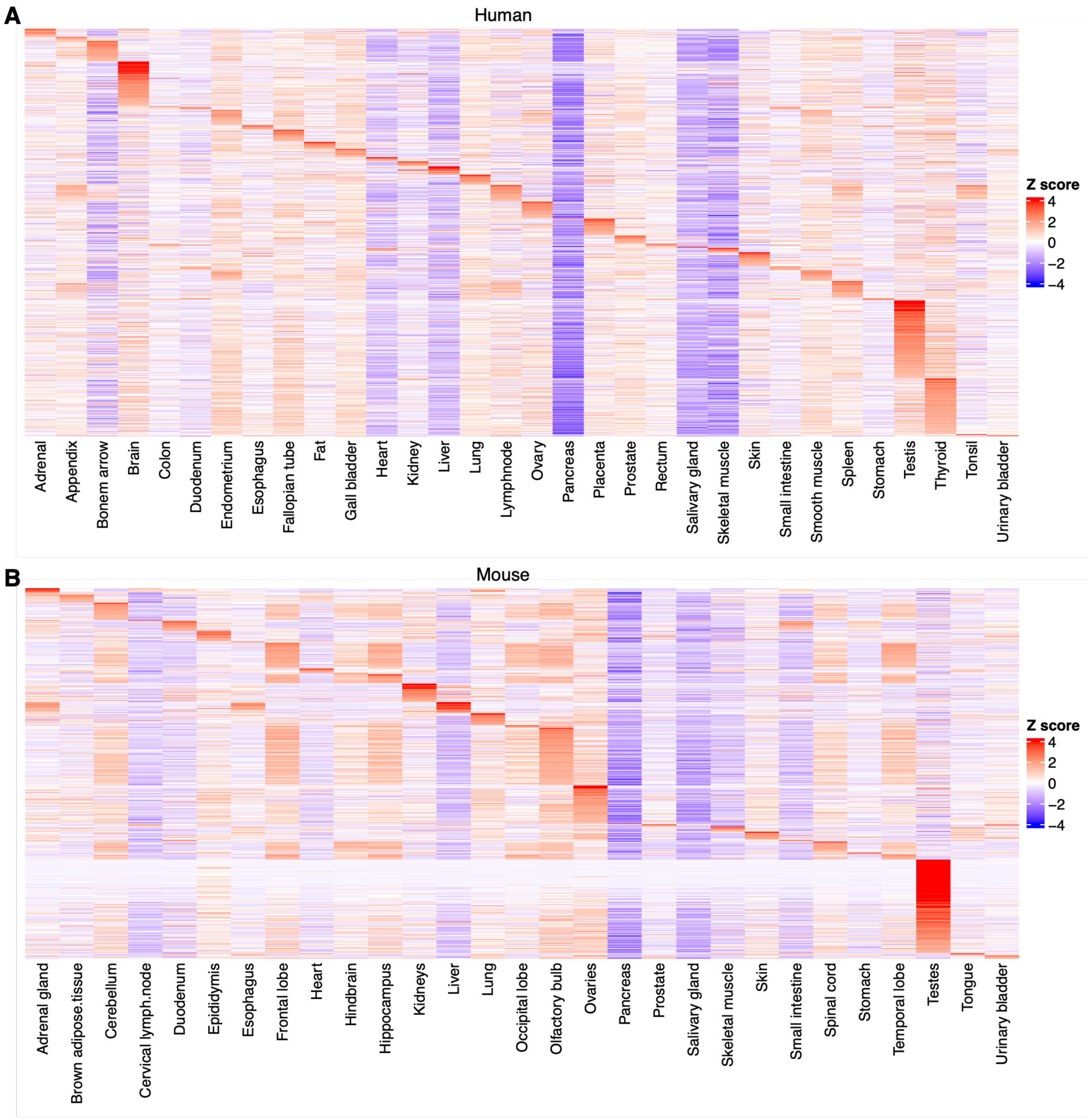


**Supplementary figure 14.** Heatmap showing the transcription levels of ncORF-containing genes in humans (A) and mice (B). Only genes expressed with TPM ≥ 1 in at least one sample are shown. TPM values were normalized across samples by Z-score transformation prior to visualization. ncORF, non-canonical open reading frame; TPM, transcripts per million.

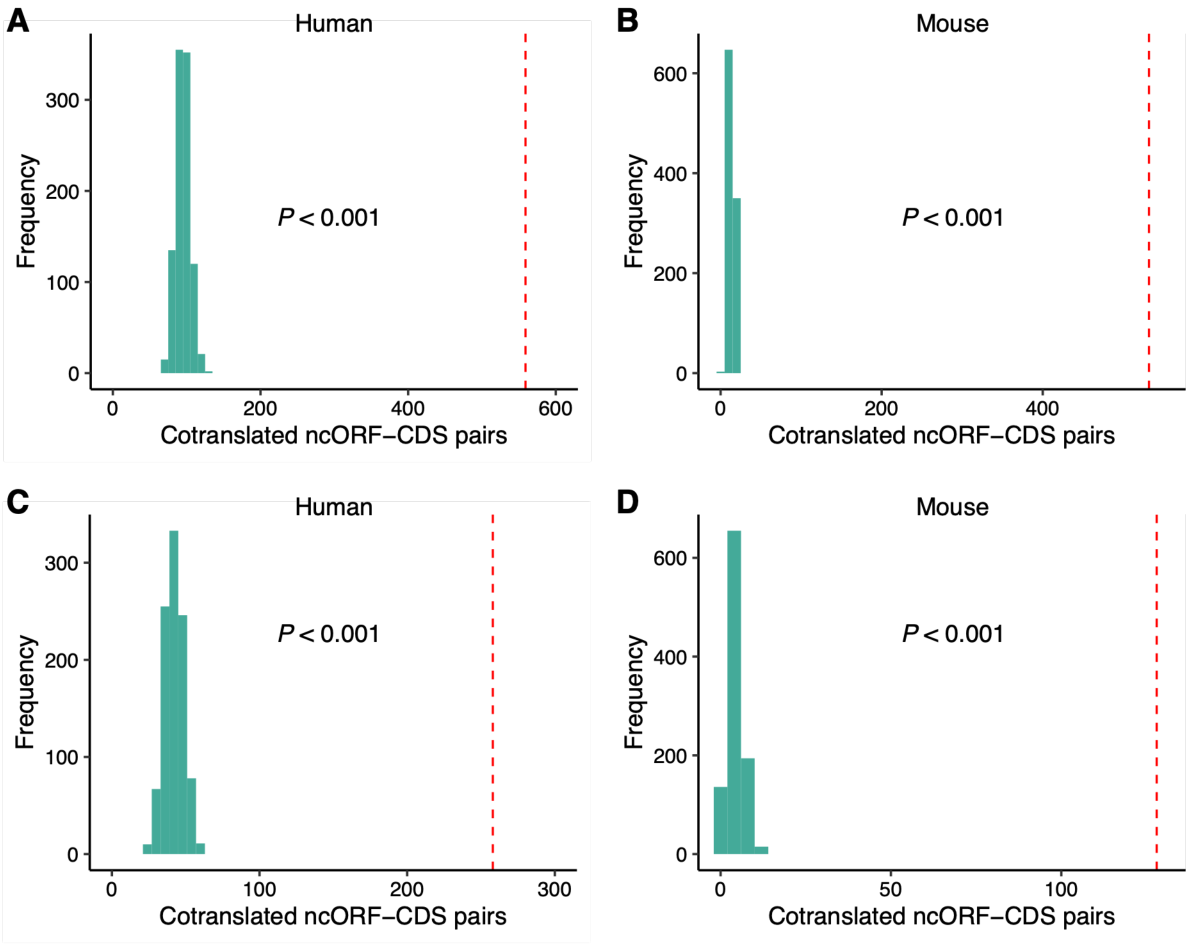


**Supplementary figure 15.** Permutation tests assessing co-translation between ncORFs and main CDSs within the same genes. (**A-B**) Distribution of the number of ncORFs that co-translate with corresponding main CDSs of the same genes in humans (A) and mice (B) based on the permutation analysis. Red dashed lines mark the observed counts. Empirical *P* values were indicated. (**C-D**) Same analyses as in (A) and (B), but excluding ncORFs overlapping with any CDS regions. CDS, coding sequence; ncORF, non-canonical open reading frame.

### Supplementary Tables

**Supplementary table 1.** High-quality ribosome profiling libraries used for non-canonical open reading frame annotation.

**Supplementary table 2.** Clusters of genomically overlapping GENCODE non-canonical open reading frames.

**Supplementary table 3.** List of non-canonical open reading frames annotated in this study.

**Supplementary table 4.** Known non-canonical open reading frames previously reported to encode functional microproteins and independently rediscovered here.

**Supplementary table 5.** Non-canonical open reading frames containing Pfam domains.

**Supplementary table 6.** Evolutionary characteristics and constraints of non-canonical open reading frames.

**Supplementary table 7.** List of ribosome profiling libraries used for translation quantification.

### Supplementary References

1. Lei, T., Chang, Y., Yao, C., and Zhang, H. (2023). A systematic evaluation of computational methods for predicting translated non-canonical ORFs from ribosome profiling data. Journal of Genetics and Genomics *51*, 105-108. 10.1016/j.jgg.2023.08.010.
